# Deletion of hypoxia-inducible factor prolyl 4-hydroxylase 2 in *FoxD1*-lineage mesenchymal cells leads to congenital truncal alopecia

**DOI:** 10.1101/2021.03.31.437126

**Authors:** Ann-Helen Rosendahl, Mia Monnius, Anu Laitala, Antti Railo, Ilkka Miinalainen, Ritva Heljasvaara, Joni M. Mäki, Johanna Myllyharju

## Abstract

Hypoxia-inducible factors (HIFs) induce hundreds of genes regulating oxygen homeostasis in tissues. Oxygen sensors of the cells, the HIF prolyl 4-hydroxylases (HIF-P4Hs), regulate the stability and activity of HIFs in an oxygen-dependent manner. In this study, we show that lack of *Hif-p4h-2* in *FoxD1*-lineage mesodermal cells interferes the normal development of hair follicles (HF) in mice. The *FoxD1*-lineage cells were found to be mainly mesenchymal cells located in the dermis of truncal skin, including the cells composing the dermal papilla of the HF. Upon *Hif-p4h-2* inactivation, HF development was disturbed during the first catagen leading to formation of large epithelial lined HF cysts filled by unorganized keratins, which eventually manifested as truncal alopecia. The depletion of *Hif-p4h-2* led to HIF stabilization and dysregulation of multiple genes involved in keratin formation, HF differentiation, and HIF, TGFβ and Notch signaling. The failure of the controlled process of HF cycling is likely to be mechanistically caused by disruption of the precise and timely interplay of the HIF, TGFβ and Notch pathways. In summary, we show here for the first time that HIF-P4H-2 function in *FoxD1*-lineage cells is essential for the normal development and homeostasis of HFs.

## INTRODUCTION

Cells have an intrinsic response to low O_2_ concentrations that is controlled by the hypoxia-inducible transcription factor (HIF, αβ dimers) (Myllyharju 2013). The HIFα subunits are negatively regulated by the HIF prolyl 4-hydroxylases 1-3 (HIF-P4Hs 1-3, also known as PHD 1-3 and EGLN2, 1 and 3, respectively) in normoxia (Bruick and McKnight 2001, Epstein *et al*. 2001, Ivan *et al*. 2001 and 2002, Jaakkola *et al*. 2001), HIF-P4H-2 being the main isoform regulating HIFα availability (Berra *et al*. 2003). In normoxia the HIF-P4Hs hydroxylate HIFα targeting it for proteasomal degradation, while in hypoxia the O_2_-dependent HIF-P4Hs are inhibited, which leads to accumulation of HIFα, and induction of the hypoxia response pathway by upregulation of HIF target genes (Myllyharju 2013, Schofield and Ratcliffe 2004).

Physiological hypoxia is vital for morphogenesis of tissues (Dunwoodie 2009, Giaccia *et al*. 2004). In mature skin partial O_2_ pressure (pO_2_) ranges between 0.2-10% and in hair follicles pO_2_ is 0.1-0.8% (Rezvani *et al*. 2011). HIF1α protein is abundant in hair follicles (HFs) and sebaceous glands, and is present at low levels in the basal keratinocyte layer (Cowburn *et al*. 2014, Evans *et al*. 2006, Rosenberger *et al*. 2007). HIF1α enhances keratinocyte and dermal fibroblast mobility and promotes cell proliferation and survival (Cowburn *et al*. 2014). HIF2α is sporadically seen in the dermal area of the skin (Rosenberger *et al*. 2007). It is also localized in the HF bulb precortex area above the matrix cells and is involved in the hair production and cell differentiation in HFs (Imamura *et al*. 2014).

In mammals, HFs have an ability to regenerate in cycles of growth (anagen), involution (catagen) and rest (telogen). This regenerative potential is plausible because of a reservoir of multipotent stem cells (SCs) located in the HF bulge area (Cotsarelis *et al*. 1990). Morphogenesis of mouse HFs starts at embryonic day 12.5 (E12.5) with the patterning of perfectly ordered pre-germ patches in the epidermis (Paus *et al*. 1999). At E14.5 epidermal signaling from the epidermal thickening (i.e. placode) forces mesenchymal fibroblasts to accumulate underneath the placodes and to form a dermal condensate that will later differentiate into dermal papilla (DP) cells of the mature HF that forms around two weeks after birth. The HF morphogenesis can be seen as an anagen-like growth phase, after which the HF proceeds to the catagen-phase of its first HF cycle (Driskell *et al*. 2011, Fuchs 2007, Krause and Foitzik 2006).

HF SCs express hypoxia-inducible genes (Rathman-Josserand *et al*. 2013), but the role of hypoxia signaling in HF development and homeostasis has not been elucidated. In this study we inactivated the main HIF regulator *Hif-p4h-2* in Forkhead box D1 (*FoxD1*)-lineage mesenchymal cells in mice. This resulted in postnatal truncal alopecia, with a premature catagen initiation and epidermal cyst formation. We show here for the first time that HIF-P4H-2 regulation of hypoxia signaling is crucial for the development and homeostasis of the hair and HF cycling.

## RESULTS

### Inactivation of *Hif-p4h-2* in *FoxD1*-lineage dermal cells results in truncal congenital alopecia

A conditional *Hif-p4h-2* (*Hif-p4h-2*^*loxP/loxP*^) mouse line was generated as described in Materials and Methods. To examine the role of HIF-P4H-2 in the dermal *FoxD1*-lineage cells we crossbred transgenic *FoxD1-Cre* (*FoxD1*^*Cre/+*^) mice with *Hif-p4h-2*^*loxP/loxP*^ mice to obtain conditional *Hif-p4h-2*^*loxP/loxP*^*;FoxD1*^*Cre/+*^ knock-out (cKO) mice (Supplemental Fig. S1a-f). Truncal hair was lost in the cKO mice and their size was smaller (Fig. 1A, B). Cranial hair of the cKO mice was normal and some hair existed around the ankles, around the tail base and sparsely on the abdominal side of the body (Fig. 1A). Formation of whiskers, nails and teeth was apparently normal in the cKO mice. None of the other littermate genotypes obtained from the matings displayed the alopecia phenotype and were used as littermate controls in the following analyses. Besides alopecia the cKO animals developed polycythemia (Supplemental Fig. S1G-I), as described previously (Kobayshi *et al*. 2016).

**Figure 1.**
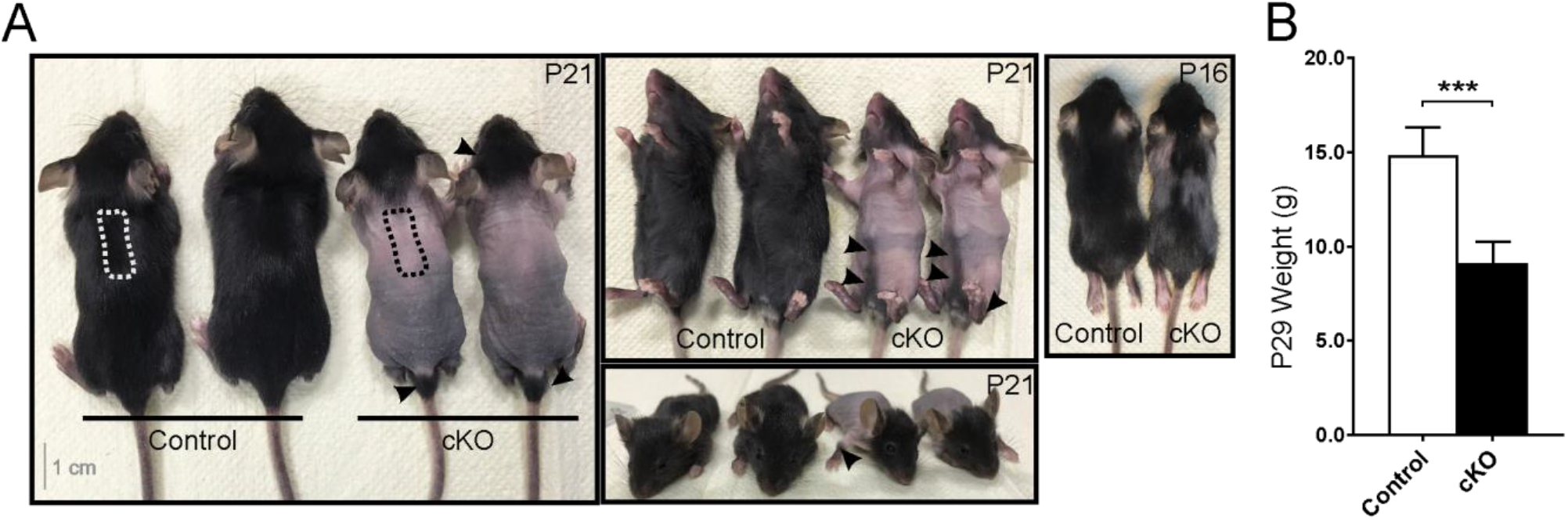
Inactivation of *Hif-p4h-2 in FoxD1*-lineage dermal cells causes congenital truncal alopecia. (A) The cKO mice have progressive congenital truncal alopecia, but maintain normal hair on the head and neck and have also traces of hair around the ankles, tail base, and on the abdominal side of the body (arrowheads). Skin biopsies for further analyses were taken from the upper back skin region indicated with a dashed line. (B) Weight difference between control (n = 6) and cKO (n = 4) mice at P29. Data are presented as mean ± S.D. *** P<0.001.

### Inactivation of *Hif-p4h-2* in *FoxD1*-lineage cells leads to disruption of HF cycling and formation of epidermal cysts

HF morphogenesis at E14.5-P14 occurred normally in the cKO mice (Fig. 2A). First differences were observed at P15 in the cKO HFs as a failure to maintain the integrity of the upper permanent part of the HF (Fig. 2A). Subsequently, at P16 the HF bulb of the cKO mice had progressed to a late stage catagen, while the control mouse HF reached a similar stage two days later (Fig. 2A). As the HF cycle progressed, the superficial part of the HFs in the cKO mice failed to maintain its structure and rigidity. The hair formation failed, and the infundibulum and isthmus of the HF expanded to an epidermal cyst filled with keratin and hair fragments (Fig. 2A, Supplemental Fig. S2). At P24 the first anagen started normally in the control mice by creating a new HF that engulfs the DP cells, while the club hair ensuring coating at all times remained from the morphogenesis and rested in its own silo in the HF upper part (Fig. 2A) (Higgins *et al*. 2009). Neither club hair formation nor initiation of a new anagen phase was observed in the P24 HFs of the cKO mice (Fig. 2A). From the beginning of telogen of the first HF cycle (P21), the number of normal HFs in the cKO mice was significantly reduced and epidermal cysts were prevalent (Fig. 2A-C, Supplemental Fig. S2). The cysts were located in the upper part of the HF (Fig. 2A), mainly comprising infundibulum and isthmus, which are the permanent parts of the HF, and not involved in the HF cycling.

**Figure 2.**
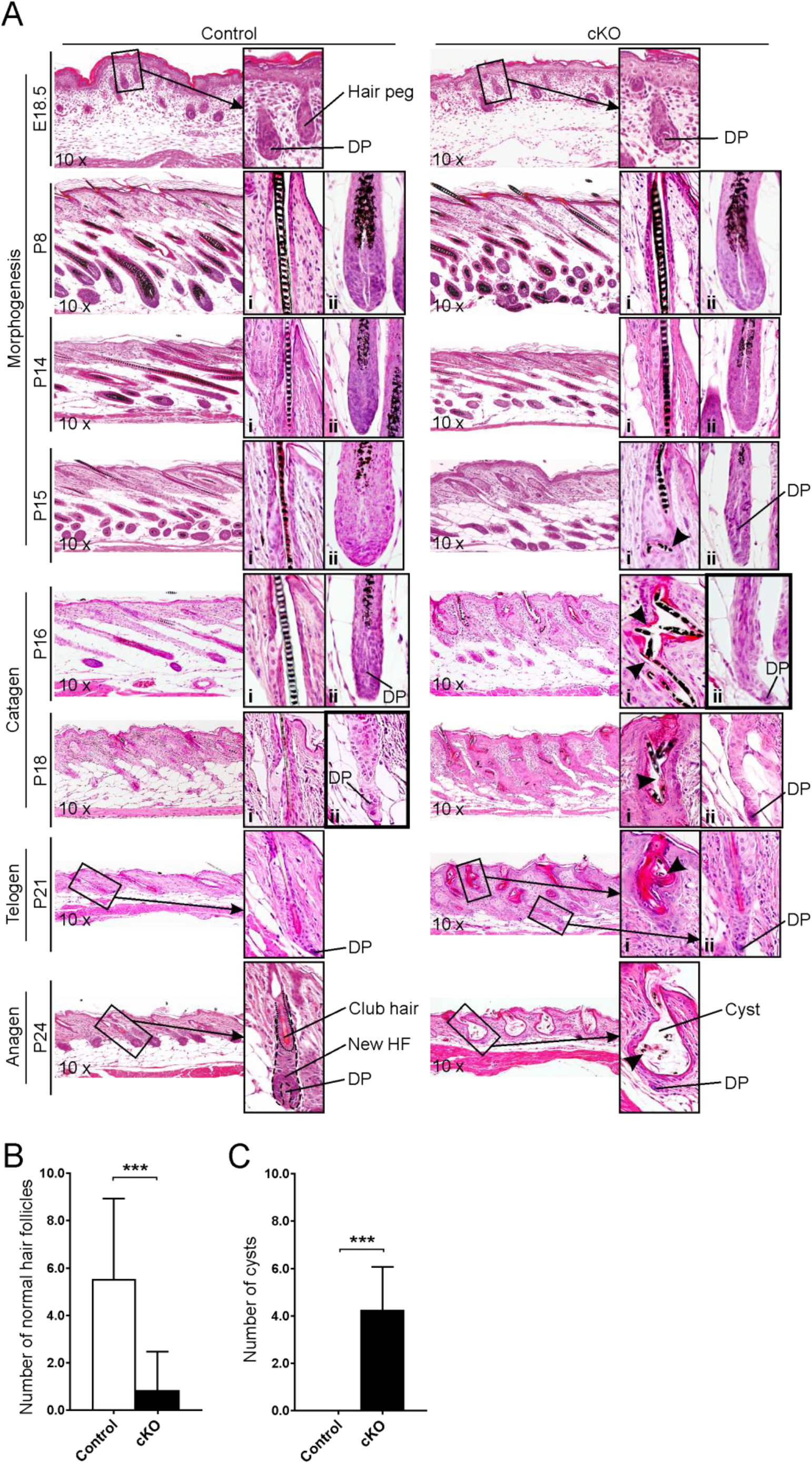
Inactivation of *Hif-p4h-2* in *FoxD1-*lineage dermal cells disturbs skin and HF homeostasis and causes epidermal cyst formation and premature catagen induction. (A) H&E staining of dorsal skin of control and cKO mice at indicated time points. Normal HF stages are indicated on the left side of the images. The first magnified insert (i) shows the infundibulum and isthmus area of the HF, while the second insert (ii) shows the bulb area of the HF. Breaking of the hair shaft to small pieces is indicated with arrowheads in the cKO mice. A late stage catagen-phase is indicated in the HF bulb inserts (ii) with a thicker frame. At the initiation of a new anagen-phase at P24 the control HF is showing one club hair and the beginning of a new HF, which is not occurring in the cKO mice. (B) Number of normal HFs per 4 visual fields/mouse in control (n = 15) and cKO (n = 18) mice (P21, P27, P29, P33). (c) Number of cysts per 4 visual fields/mouse in control (n = 37) and cKO (n = 35) mice (P21, P24, P27, P29, P33). Data are presented as mean ± S.D. * P<0.05; *** P<0.001.

### *FoxD1*-lineage cells are located in the truncal dermis and DP of the HF

To identify the distribution of the *FoxD1*-lineage mesenchymal cells in the skin we crossbred the *FoxD1*^*Cre/+*^ mice with double fluorescent *Cre* reporter mice, the *Rosa26*^*mT/mG*^ mice. *FoxD1*-*Cre* mediated deletion was observed in dermal cells with a fibroblast-like morphology, as well as in the DP cells in the HFs at P14 (morphogenesis), P21 (1^st^ telogen) and P27 (early 1^st^ anagen) (Supplemental Fig. S3). In the cranial P21 skin sections only a few *FoxD1*^+^ cells were observed in the dermis and no *FoxD1*^+^ DP cells were detected in the HF (Supplemental Fig. S3). These observations are supported by studies showing that the cells in head and neck dermis originate from the neural crest of the ectoderm (Couly and Le Douarin 1988), while the cells in ventrolateral and dorsal dermis are of mesodermal origin (Olivera-Martinez *et al*. 2004, Driskell *et al*. 2011). We also confirmed the localization of *FoxD1*-lineage cells in the cKO mice by producing cKO mice that express simultaneously the *Rosa26*^*mT/mG*^ and *FoxD1-Cre* transgenes. The localization of *FoxD1*^*+*^ cells was not affected by *Hif-p4h-2* deletion (Supplemental Fig. S3). The difference in the origin and hence *FoxD1* expression in the cranial and truncal dermal fibroblast-like cells thus explains the distribution of the alopecia in the cKO animals and underlies the importance of HIF-P4H-2 in hair development.

### Inactivation of *Hif-p4h-2* in *FoxD1*-lineage cells leads to activation of the hypoxia response pathway in the skin

To confirm the deletion of *Hif-p4h-2* in dermal *FoxD1*-lineage cells, we analyzed the protein expression of HIF-P4H-2, HIF1α and HIF2α from the cKO and control mouse skin (P16). Deletion of the *Hif-p4h-2* gene in the skin is solely present in the mesenchymal cells derived from the *FoxD1*-lineage, which represent only a fraction of the total cell amount in the skin (Supplemental Fig. S3). Nevertheless, a decreased amount of HIF-P4H-2 protein was observed in cKO skin *in toto* (Supplemental Fig. S1E, F) and HIF1α and HIF2α were markedly stabilized in the cKO skin (Fig. 3A). Furthermore, a systematic mRNA upregulation of known HIF target genes was observed in the cKO skin from P14-16 onwards (Fig. 3B-F), while no difference was observed in the mRNA level of a non-target gene *Hif-p4h-1* (Fig. 3G).

**Figure 3.**
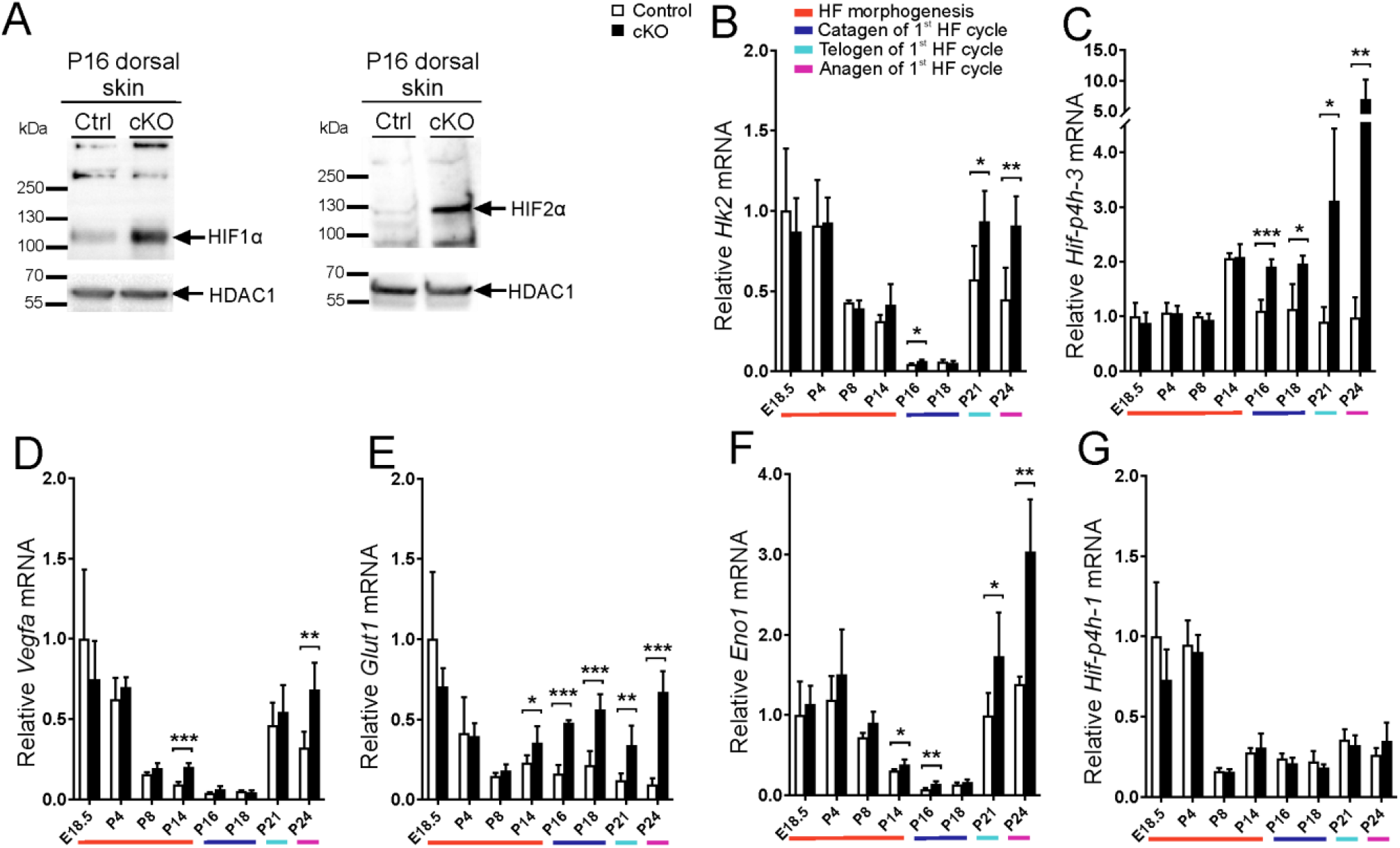
HIF1α and HIF2α are stabilized and expression of HIF target genes is upregulated in *cKO* skin. (A) Western blot analysis of HIF1α and HIF2α in control and *cKO* P16 skin. Histone deacetylase 1 (HDAC1) was used as a loading control. (B-G) qPCR analysis of mRNA expression of HIF target genes *Hk2* (B), *Hif-p4h-3* (C), *Vegfa* (D), *Glut1* (E), *Eno1* (F), and the non-HIF target *Hif-p4h-1* (G) at indicated time points. The colours beneath the bar charts indicate the HF cycle stages, n = 4-7 per genotype. Data are presented as mean ± S.D. * P<0.05; ** P<0.01; *** P<0.001.

### Inactivation of *Hif-p4h-2* in *FoxD1*-lineage cells leads to upregulation of genes involved in skin barrier function and hair formation

We next analyzed the cKO skin structure and epidermal cyst formation in more detail by scanning electron microscopy, which showed that the skin had an abnormal appearance, with sparse and irregular hair formation and atypical keratin shedding from the skin surface (Fig. 4A). The cysts in the cKO skin contained accumulated keratin and small pieces of broken hair shafts and were typically located in the upper part of the dermis. Only a few fragile and irregular hair shafts existed in the cKO skin (Fig. 4A). We next studied the expression and distribution of various keratins and keratin-related proteins in cKO skin. Proliferative progenitor cells in the epidermal basal layer and the outer root sheath (ORS) are rich in dimerized keratin 5 (KRT5) and 14 (KRT14) (Botchkarev and Paus 2003, Hsu *et al*. 2014a, Moll *et al*. 2008), which provide mechanical support and cytoprotection in the basal cells (Alam *et al*. 2011). The basal cells are responsible for the integrity of the epidermal basal layer and are the origin of the non-proliferative upper layers of the skin (Hsu *et al*. 2014a). The spinous and granular cell layers of the epidermis express dimerized KRT1 and KRT10, which have a major role in epidermal barrier formation (Botchkarev and Paus 2003, Roth *et al*. 2012). As the keratinocytes differentiate to a post mitotic stage and migrate towards the skin surface, *Krt5*/*Krt14* expression is reduced and *Krt1*/*Krt10* expression is induced (Alam *et al*. 2011). The cysts in the cKO skin were filled with KRT5 (Fig. 4B), and its staining in the skin was diffuse in the epidermis and cyst edge, whereas in the controls the staining was specific to the epidermal basal cells and to the HF (Fig. 4B). *Krt5* expression was significantly increased in the cKO mice at the end of morphogenesis (P14) and at the beginning of anagen of the first HF cycle (P24), but was unaffected in other time points (Fig. 4C). *Krt14* mRNA level was significantly upregulated in cKO mouse skin starting from P16, the higher expression level persisting during the remaining HF cycle (Fig. 4C). The upper epidermal layer was intensely stained with KRT1 in the cKO skin (Fig. 4B). KRT1 was localized on the edges of the cysts, suggesting that the cKO mice attempt to develop the inner root sheath (IRS) of the HFs (Fig. 4B). In cKO mice a premature induction of *Krt1* expression was observed at P14 and it persisted throughout the cyst formation, while less pronounced effects on *Krt10* expression were observed at P14-18 (Fig. 4C).

**Figure 4.**
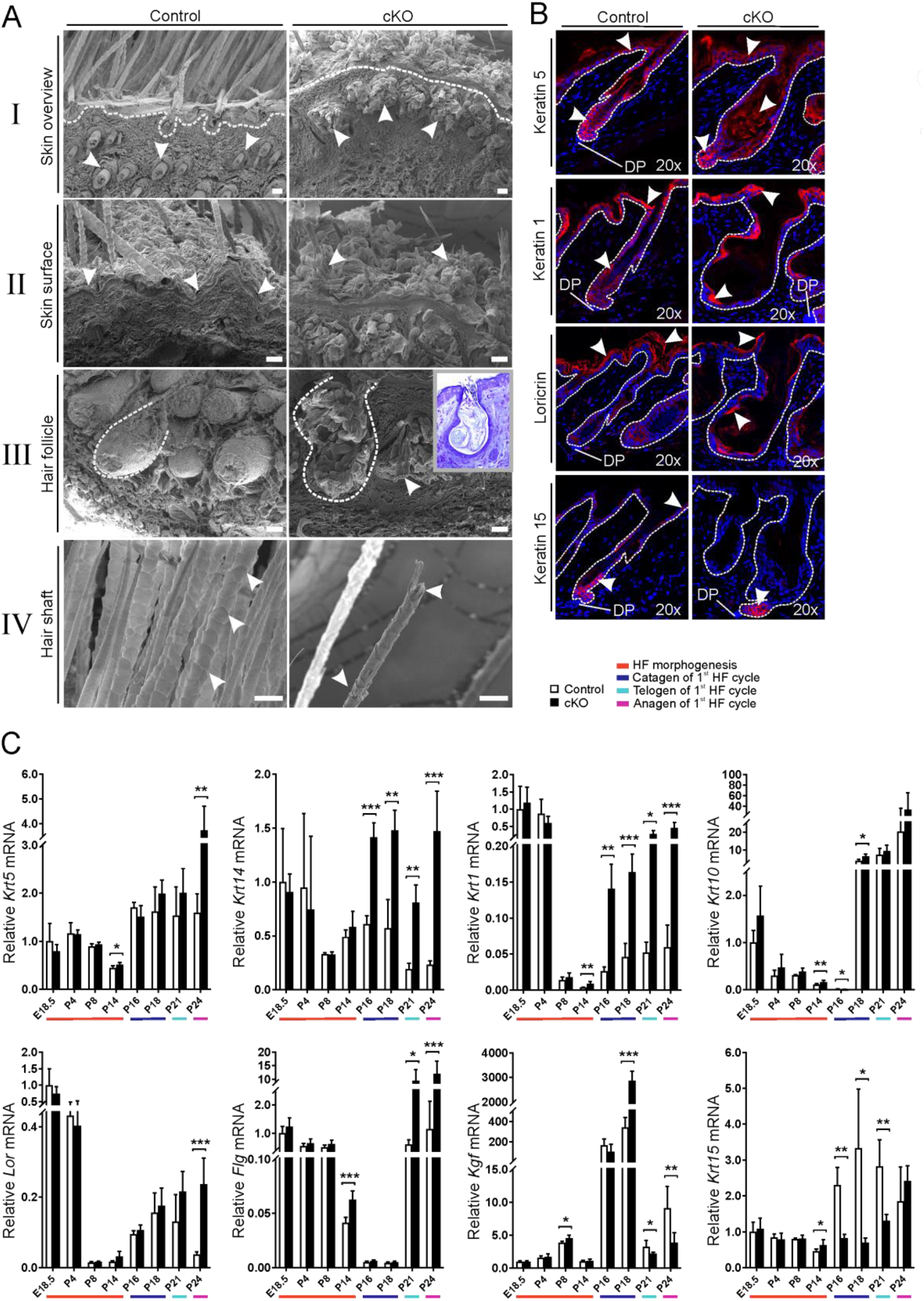
Inactivation of *Hif-p4h-2* in *FoxD1*-lineage dermal cells increases the expression of keratins and changes their distribution in the skin. (A) Scanning electron microscopy analysis of P24 control and cKO skin. I: HFs are indicated with arrowheads and the basement membrane is visualized with a dashed line. In cKO skin keratin-filled cysts mainly located in the upper part of the dermis are observed. II: Keratins on the skin surface are indicated with arrowheads. They separate from the skin in control mice as sheaths, while they emerge as clusters on the cKO skin. III: HFs are indicated with a dashed line. An arrowhead indicates a fragment of a hair shaft inside a cyst in cKO skin. The cKO insert shows the histological appearance of a similar cyst. IV: the hair shaft scaling is indicated with arrowheads. The cKO skin hair shaft scaling is irregular, and the hair shaft is fragile and breaks easily. Scale bars: 20 µm. (B) Immunofluorescence staining of keratins 1, 5 and 15, and loricrin in control and cKO dorsal skin at P21. Examples of positive signals are indicated by arrowheads. Dermal papilla (DP) is shown in the figures and the basement membrane is visualized with a dashed line. (C) qPCR analysis of mRNA expression of keratins (*Krt5, Krt14, Krt1, Krt10* and *Krt15*), loricrin (*Lor*), filaggrin (*Flg*) and keratinocyte growth factor (*Kgf*) in control and cKO mouse skin. The colours beneath the bar charts indicate the HF cycle stages, n = 4-7 mice per genotype. Data are presented as mean ± S.D. * P<0.05; ** P<0.01; *** P<0.001.

Loricrin (LOR) is expressed in the IRS and *stratum corneum* and contributes to the protective barrier of the cornified envelope (Botchkarev and Paus 2003). The apical side of the cysts of the cKO mice produced LOR (Fig. 4B). LOR was also localized on the outermost surface of the skin in both genotypes (Fig. 4B). No differences were observed in *Lor* mRNA expression at E18.5-P21, but at the induction of anagen of the first HF cycle (P24) *Lor* expression was maintained at a higher level in the cKO (Fig. 4C). Filaggrin (FLG) is necessary for the formation and continuance of the cornified envelope by binding to keratins, but also for the flattening of the cells in the cornified layer (Rundle *et al*. 2017). *Flg* mRNA expression was higher in the cKO at P14, normalized during catagen, and enhanced again during telogen (P21) and beginning of anagen (P24) of the first HF cycle, relative to control (Fig. 4C).

### Inactivation of *Hif-p4h-2* in *FoxD1*-lineage cells leads to abnormal expression of keratinocyte growth factor and keratin 15 and disturbs cornification of the inner root sheath

DP cells influence the HF bulge SCs to progress to the next phase of the HF cycle (Hsu *et al*. 2014a). We examined whether inactivation of *Hif-p4h-2* in the DP cells affects their communication with the bulge SCs, which could result in the interruption of the development of the HF and subsequent cyst formation. Hypoxia-inducible keratinocyte growth factor (KGF) is produced by mesenchymal fibroblasts to ensure the colony formation and proliferation of the overlying keratinocytes. The DP cells produce KGF especially during the initiation of anagen to influence the bulge SCs to create a new HF (Guo *et al*. 1993, Hsu *et al*. 2014a, Tsuji *et al*. 2014). At the end of the catagen (P18) a strong upregulation of *Kgf* expression was observed in the cKO mice relative to control, which later at P21 and P24 converted to a marked downregulation of the *Kgf* mRNA (Fig. 4C). This indicates that the initiation of the anagen phase by the cKO DP cells in the cyst bulge SCs is disturbed.

The bulge SC marker KRT15 is co-produced with the KRT5/KRT14 heterodimer in the basal layers of the epidermis (Bose *et al*. 2013). In cKO P21 HFs KRT15^+^ cells (i.e. potential SCs) were located specifically in the bulb area of the cyst, while in the controls faint staining of KRT15^+^ cells was detected within the basement membrane of the epidermis and in the bulge (Fig. 4B). In some of the cKO hair cysts KRT15^+^ staining was completely absent, while in other cysts smaller clusters of KRT15^+^ staining was observed. The *Krt15* mRNA level was significantly higher at P14 in the cKO skin, while during the HF catagen and telogen (P16-P21) the level was significantly reduced relative to controls (Fig. 4C). This data suggests that the bulge SC population is abnormally distributed in the cKO mice, causing interference in the signaling from the DP cells to the bulge SCs.

Transmission electron microscopy revealed that formation of both the hair shaft and the HF was disturbed in the cKO mice. In a cross-section of control P14 HF and hair shaft all cell layers were visible (Fig. 5A). In the cKO P14 HF and hair shaft the same cell layers were present, but they had morphological discrepancies. The Henle’s layer in the cKO IRS was not as uniform in thickness as in the control and the cKO ORS layers contained unidentified loose material in the cytoplasm. The cells also seemed to have lost their polarity and normal morphology. The P24 cKO cysts were composed of an unstructured mass of hair shaft medulla, cortex and cuticle layers that were not adhering to each other normally but were peeling off from the HF structure (Fig. 5B). Moreover, the IRS of the cKO mice completely lacked the Henle’s layer (Fig. 5B), which is an important structure as it is the first of the IRS layers to cornify as the cells travel up from the bulb of the HF and mature (Joshi 2011). It moulds and protects the hair shaft and forms a cornified sheet around the HF to keep its structure intact (Alibardi and Bernd 2013).

**Figure 5.**
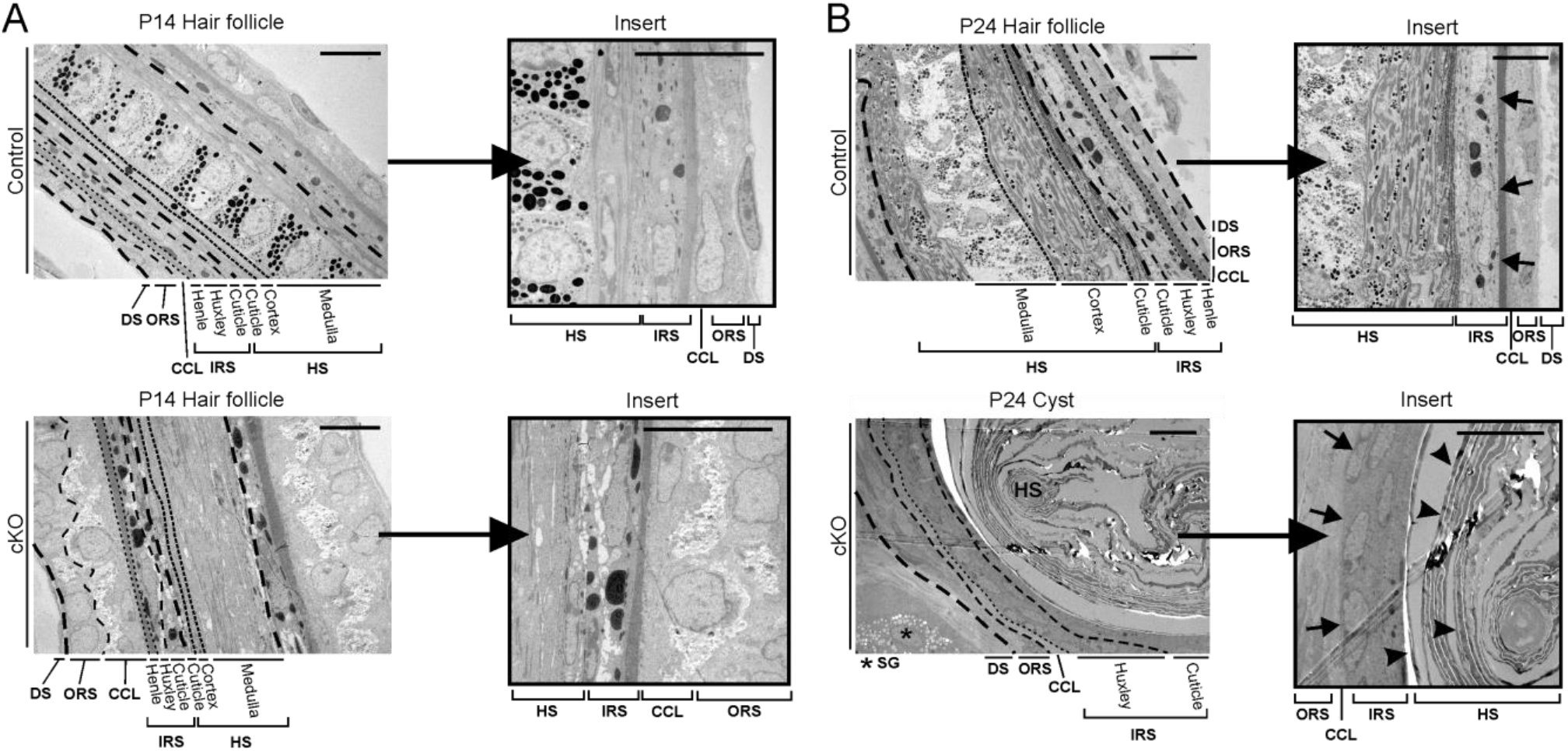
Epidermal cysts in the cKO mouse skin are poorly differentiated and do not develop Henle’s layer of the inner root sheath. Transmission electron microscopy analysis of control HF and cKO cysts at P14 (A) and P24 (B). Cell layers are separated with dashed lines. The control HF Henle’s layer is indicated in the P24 (B) magnified insert with arrows. In the P24 cKO insert the arrows indicate the place where Henle’s layer should develop. Arrowheads show keratin peeling off the sides of the cyst as layers creating the keratin mass inside the cyst. Scale bars: 10 μm. Abbreviations: hair shaft (HS), inner root sheath (IRS), companion cell layer (CCL), outer root sheath (ORS), dermal sheath (DS), Henle’s layer (He), Huxley’s layer (Hu), cuticle (Cu), cortex (Co), medulla (Me), sebaceous gland (SE).

### Inactivation of *Hif-p4h-2* in *FoxD1*-lineage cells does not affect apoptosis and proliferation of hair follicle keratinocytes

Proliferation and apoptosis of HF cells are strictly regulated during HF cycling (Alonso and Fuchs 2006, Magerl *et al*. 2001). No difference between the genotypes was observed in the proliferation of P21 HF keratinocytes (Supplemental Fig. S4A, B). Since the diameter of HFs is consistent with an increase or decrease in HF proliferation (Imamura *et al*. 2014), we also measured the diameter of the HFs but no differences were observed between the genotypes (data not shown). Analysis of apoptotic keratinocytes by TUNEL assay in P21 HFs showed no differences between the genotypes (Supplemental Fig. S4C, D). Despite this, we analyzed the expression level of the BCL2/adenovirus E1B 19kDa interacting protein 3 (*Bnip3*), a HIF target gene (Gordan and Simon 2007, Kothari *et al*. 2003) that has been linked to increased autophagocytosis besides apoptotic activity (Burton and Gibson 2009). We found a significantly increased *Bnip3* mRNA level in the cKO skin at P8, P14 and P24 (Supplemental Fig. S4E).

### Inactivation of *Hif-p4h-2* in *FoxD1*-lineage cells leads to upregulation of TGFβ signaling in hair follicles

Transforming growth factor-β (TGFβ) has an important role in HF development and cycling (Stenn and Paus 2001). TGFβ1 induces anagen by activating apoptosis and reducing keratinocyte proliferation, whereas TGFβ2 has been linked to the induction of the HF growth during morphogenesis, and both of them have been implicated in the anagen-catagen switch (Foitzik *et al*. 1999, Soma *et al*. 2002). We observed differential mRNA expression patterns of the TGFβ isoforms as well as their target genes in the cKO and control mouse skin from P14 onwards (Fig. 6A-E, Supplemental Fig. S5A-D). The mRNA expression of TGFβ1 was typically upregulated (Fig. 6SA) in most of the timepoints, while TGFβ2 was downregulated and TGFβ3 was mostly unchanged (Supplemental Fig. S5A-B). Since especially Periostin (*Postn*) and plasminogen activator inhibitor 1 (*Pai1*), target genes of TGFβ signaling, showed significant and parallel changes in their expression patterns (Fig. 6B-E, Supplemental Fig. 5C-D), we analyzed SMAD2 phosphorylation within the HF. Phosphorylation of SMAD2 was significantly higher in the cKO HF cysts at late anagen (P29) when compared to the control HFs (Fig. 6F, G), indicating upregulation of canonical TGFβ signaling in the cKO HF cysts.

**Figure 6.**
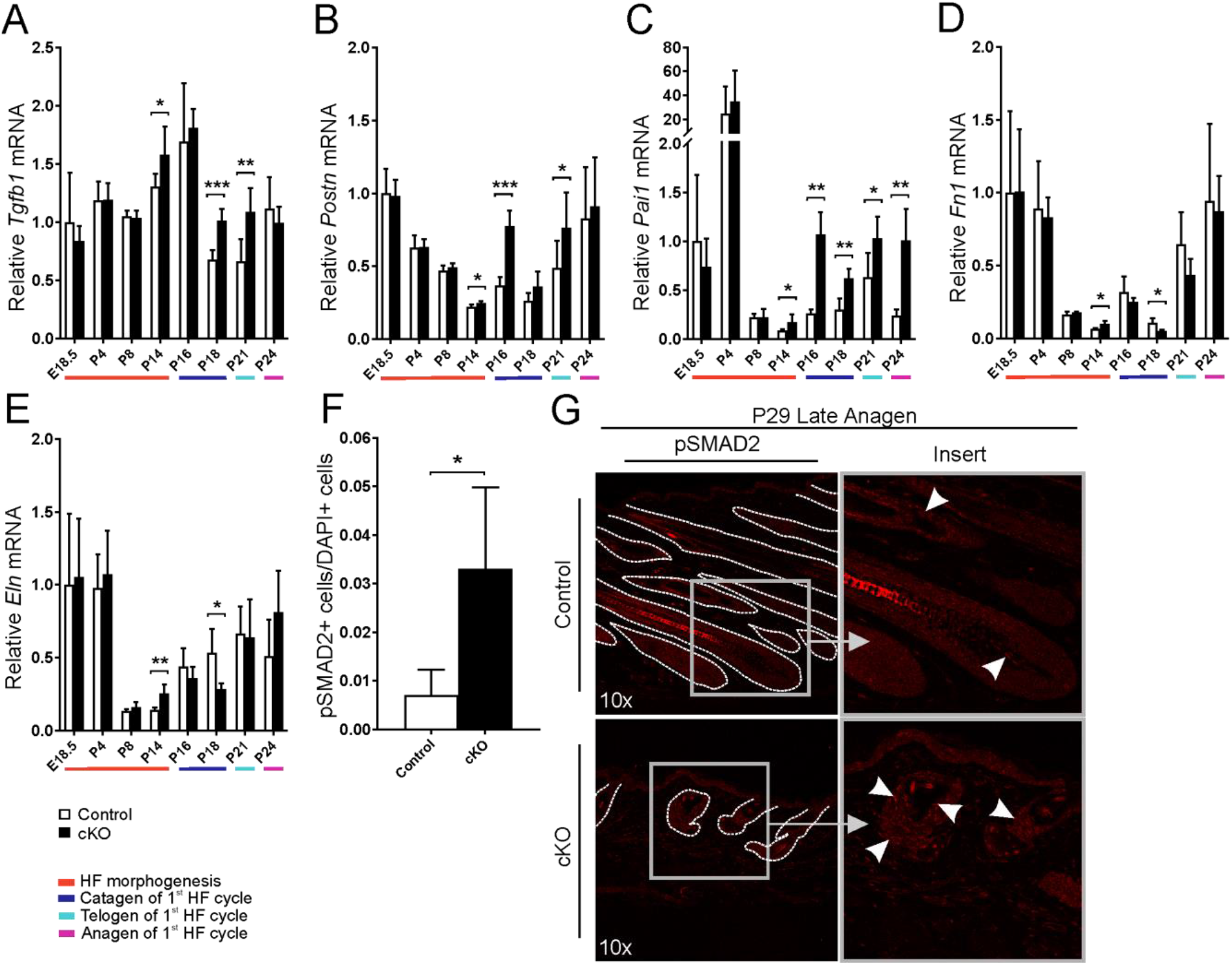
TGFβ signaling is upregulated in the cKO skin. qPCR analysis of mRNA expression of *Tgfb1* (A) and the TGFβ target genes *Postn* (B), *Pai1* (C), *Fn1* (D), and *Eln* (E) at indicated time points. The colours beneath the bar charts indicate the HF cycle stages, n = 4-7 per genotype. (F) Morphometric analysis of pSMAD2^+^ keratinocytes in the HF. Control (n = 5), cKO (n = 6). Data are presented as mean ± S.D. * P<0.05; ** P<0.01; *** P<0.001. (g) Representative images of immunostaining of pSMAD2 in the skin. Arrowheads indicate pSMAD2 staining in the magnified inserts. HF structures in the control and cysts in the cKO mouse skin are indicated with a dashed line.

### Inactivation of *Hif-p4h-2* in *FoxD1*-lineage cells disturbs Notch signaling in the skin

NOTCH is an important factor for postnatal HF development and homeostasis (Vauclair *et al*. 2005). NOTCH is expressed abundantly both in the HF and the epidermis together with its activating ligands (Nowell and Radtke 2013, Watt *et al*. 2008). The NOTCH receptor ligands, such as delta-like ligands (DLL1, DLL3, DLL4) and Jagged ligands (JAG1, JAG2) can activate Notch signaling by ligand-receptor interaction and are expressed in NOTCH receptor neighbouring cells (Nowell and Radtke 2013). Furthermore, ADAM metalloproteinases, i.e. ADAM8, ADAM10 and ADAM17, can activate Notch signaling by extracellular cleavage, while γ-secretase releases the NOTCH intracellular domain (NICD, the activated NOTCH) in the cells, which can subsequently activate NOTCH target genes (Brou *et al*. 2000, Weber *et al*. 2011). We therefore analyzed the mRNA levels of the NOTCH receptors 1-4 (*Notch1-4*), their ligands and other activators such as selected ADAMs, and NOTCH target genes from the skin samples. We found fluctuating differences in the relative expression levels of these genes between the genotypes mostly starting around P14 (Fig. 7A-L, Supplemental Fig. S5E-I). At the protein level, the amount of NOTCH1 and its activated form NICD, as well as its target HES1 and potential activator ADAM10, were clearly increased in the cKO mouse skin at P21 (Fig. 7M). Interestingly, factor inhibiting HIF (FIH) is known to hydroxylate NICD and inhibit its function (Andersson *et al*. 2011). *Fih* mRNA level was similar between the genotypes until P24, when its mRNA level was significantly higher in the cKO mouse skin (Supplemental Fig. S5J). Interestingly, higher FIH protein amount was seen in cKO skin already at P14 (Supplemental Fig. S5K). Taken together, the data suggest that Notch signaling homeostasis is disturbed in the cKO mice.

**Figure 7.**
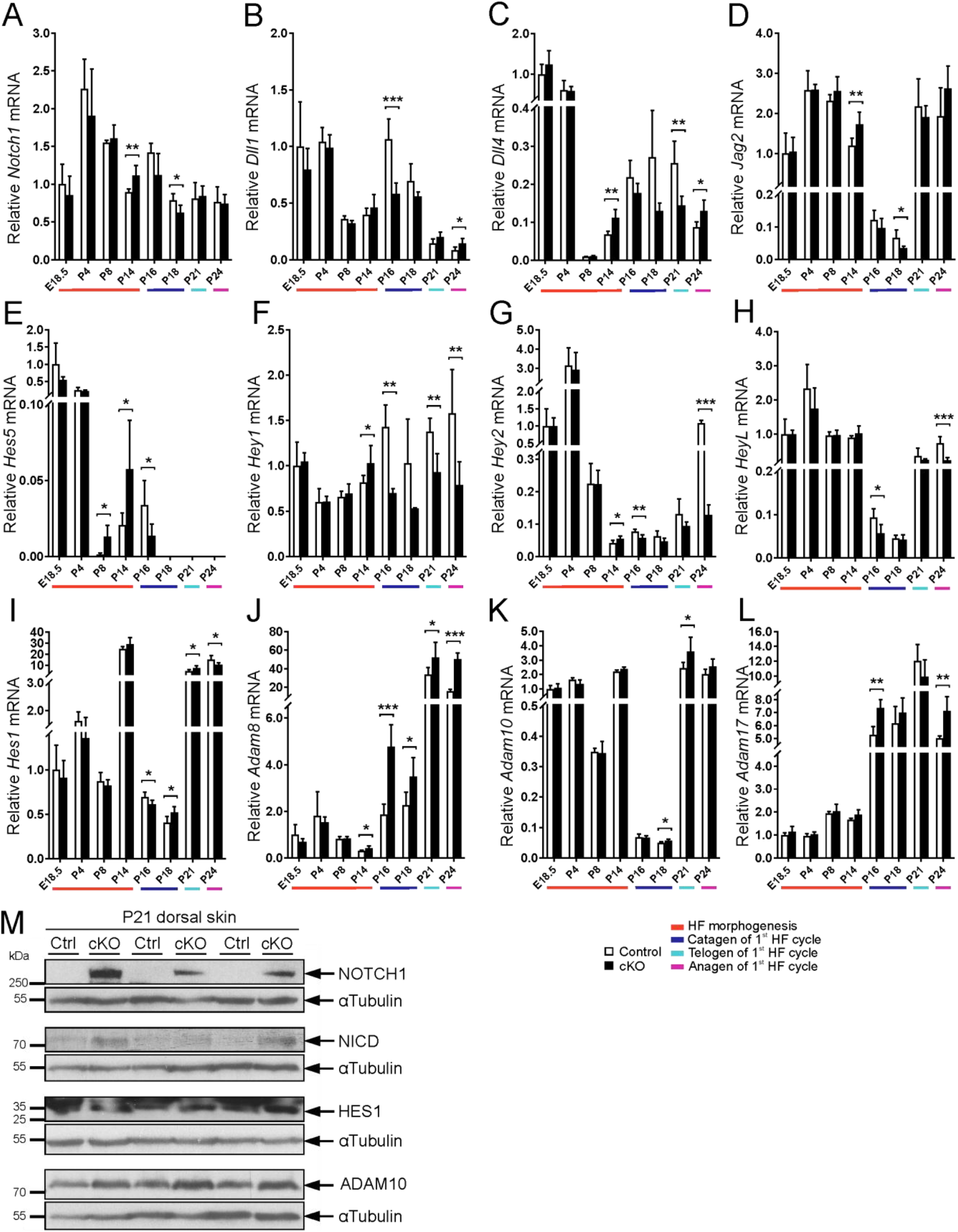
Notch signaling is distured in the cKO skin. qPCR analysis of mRNA expression of *Notch1* (A); NOTCH ligands *Dll1, Dll4*, and *Jag2* (B-D); NOTCH target genes *Hes5, Hey1, Hey2, HeyL and Hes1* (E-I); The ADAM metalloproteinases *Adam8, Adam10* and *Adam17* (J-L) at indicated time points. The colours beneath the bar charts indicate the HF cycle stages, n = 4-7 per genotype. (m) Western blot analysis of NOTCH1, NICD, HES1, and ADAM10 protein in P21 dorsal skin samples. Data are presented as mean ± S.D. * P<0.05; ** P<0.01; *** P<0.001.

## DISCUSSION

We show here for the first time that inactivation of the main oxygen sensor HIF-P4H-2 in *FoxD1*-lineage cells disrupts normal HF development and cycling. Global inactivation of HIF-P4H-2 in mouse leads to death of the embryos at E12.5-14.5 (Takeda *et al*. 2006), whereas global conditional inactivation of HIF-P4H-2 by tamoxifen administration 17.5 days after coitus and at 3 weeks of age leads to polycythemia and congestive heart failure (Minamishima *et al*. 2008). Polycythemia is also observed when HIF-P4H-2 is inactivated in the *FoxD1*-lineage (Kobayashi *et al*. 2016 and this study), as the erythropoietin producing kidney tubular interstitial fibroblasts are also derived from the *FoxD1*-lineage. However, alopecia has not been reported in the previous studies. A similar hairless phenotype has been shown in mice lacking ADAM10, NOTCH1, RBP-JK and γ-secretase, activators of the NOTCH signaling pathway (Pan *et al*. 2004, Vauclair *et al*. 2005, Weber *et al*. 2011). Like our cKO mice, these mice display an undisturbed early postnatal HF formation and have a premature catagen initiation and epidermal cyst formation.

In our cKO mice most of the mRNA level changes of the analyzed genes started around P14 in late morphogenesis and manifested as a progressive HF phenotype starting from P15, eventually leading to alopecia caused by a premature catagen initiation and epidermal cyst formation. The observed changes most probably result from altered communication between the mesenchymal and epithelial cells, since *Hif-p4h-2* is not deleted in keratinocytes, which are responsible for producing multiple types of keratins and keratin-related proteins needed for proper hair formation. From a mechanistic point of view, besides the HIF pathway, we found differential regulation of two major signaling pathways regulating the HF cycling, the Notch and TGFβ pathways.

Our results show that inactivation of HIF-P4H-2 changes Notch signaling in skin as shown by the differential expression of its receptors, target genes, ligands and other activating factors such as ADAMs (Fig. 7 and Supplemental Fig. S5). At protein level at P21 (1^st^ telogen) NOTCH1 is virtually absent in control skin while it is upregulated strongly in cKO skin, and upregulation of the activated NOTCH (NICD) is seen simultaneously with ADAM10 (Fig. 7M). However, this seemed not to lead to systematic induction of NOTCH target gene expression, which may indicate severe disturbance between activated NOTCH and transcriptional control of the target genes. It has been shown that HIF2α, which was stabilized in the cKO skin as a result of *Hif-p4h-2* deletion, may inhibit NICD activity by binding to its RBP-J-associated module domain (Gustafsson *et al*. 2005, Hu *et al*. 2014) and subsequently downregulate NOTCH1-dependent target genes, which is in accordance with the similar skin phenotype resulting from *Notch1* (Hu *et al*. 2014) and *Hif-p4h-2* deletion. Moreover, FIH has been shown to inhibit NICD, which could strengthen this hypothesis (Supplemental Fig. S5J, K). HIF1α, on the contrary, physically binds to NICD, causes its stabilization, and increases the NOTCH target gene expression (Gustafsson *et al*. 2005, Hu *et al*. 2014). From NOTCH receptors only NOTCH1 is expressed abundantly in the *FoxD1*-positive HF cells (including DP cells) and thus may be able to interact directly with HIF1α and HIF2α (or FIH) in DP cells. On the other hand, many Notch pathway genes are regulated by HIF (Andersson *et al*. 2011, Borggrefe *et al*. 2016, Zheng *et al*. 2017) and hence the stabilization of HIF caused by deletion of *Hif-p4h-2* may interfere Notch signaling via modifying the expression of for example *Notch3, Dll1, Dll4, Jag2*, and *Hes1*. The imbalance between HIF and Notch pathways is likely to play a crucial role in cyst formation and the failure in regeneration of HFs in the cKO skin.

NOTCH1 has been implicated to be important for the differentiation of both the HF matrix cells and the cells in the IRS - especially the Henle’s layer (Pan *et al*. 2004), that is absent from the cKO HF cysts (Fig. 5). The IRS protects the hair and is crucial for withholding the HF structure and for the hair formation itself (Schlake 2007). The Henle’s cell layer of the IRS is the first HF layer to reach full keratinization and is fundamental for maintaining the HF structure (Alibardi and Bernd 2013, Joshi 2011). In the cKO mice HF Henle’s cell layer disappears when the epidermal cyst structures develop and is no longer capable of maintaining the HF integrity (Fig. 5). Our results clearly show that expression of various keratins and keratin-related proteins, especially *Krt1, Krt14* and *Krt15*, is markedly distorted in cKO skin (Fig. 4C). Deletion of *Krt14* is known to enhance NOTCH1 signaling, and KRT14/KRT15 heterodimers are suggested to be important for promoting and maintaining cell proliferation in the epidermal basal cell layer (Alam *et al*. 2011). KRT15, the marker of the HF SCs, is not only expressed in higher amounts in cKO skin but has also changed its physical localization in the cKO HF (Fig. 4B). This is interesting since Notch signaling is also considered to be involved in determining the differentiation programs of the HF SCs (Kopan and Weintraub 1993, Watt *et al*. 2008). Major regulators of the HF cycling are KRT15^+^ SCs that are located at the lowest part of the permanent HF in the bulge region, and which can give rise to a new HF after signaling from the DP cells (Hsu *et al*. 2014b). In the cKO HF the intensity of KRT15 staining was much lower and restricted only to a small area above the DP, whereas in controls KRT15^+^ cells travelled towards the skin surface along the edges of the HF (Fig. 4B), which suggests that the SC differentiation, proliferation and/or migration may be interrupted in cKO mice.

While Notch signaling regulates skin cell fate decision in the HF bulge SCs (Yamamoto *et al*. 2003), TGFβ signaling is involved in both morphogenesis and HF cycling. TGFβ1 is especially important in the anagen-catagen transition, as well as in the telogen-anagen transition (Alonso and Fuchs 2006, Foitzik *et al*. 1999, Jamora *et al*. 2005). Hypoxia has been shown to regulate matrix proteinases and thrombospondin, whose activation may lead to the activation of TGFβ, and this phenomenon has been suggested to be HIF-mediated in hepatocytes (Roth and Copple 2015). Furthermore, HIF1α accumulation in alveolar macrophages is associated with TGFβ1-induced PAI1 production and is reversibly inhibited by HIF1α silencing (Ueno *et al*. 2011). Hypoxia also affects directly the SMAD2/3 complex by enhancing its transfer to the nucleus in human dermal fibroblasts, and subsequently drives them into transition to myofibroblasts (Zhao *et al*. 2017). TGFβ ligands are expressed in the DP cells, which in turn activate the overlying basal epithelial cell TGFβ signaling during catagen (Mesa *et al*. 2015). Increased TGFβ1 signaling has been shown to induce apoptosis in the HF (Soma *et al*. 2002). We observed significantly upregulated canonical TGFβ signaling in the cKO keratinocytes relative to the control (Fig. 6F, G), which was in accordance with upregulation of some of the TGFβ target genes in the mutant skin (Fig. 6B, C). Although no statistically significant increase in the apoptosis of skin cells was observed in the mutant mice (Suppl. Fig. 4C, D), differences in the mRNA level of the apoptosis-related *Bnip3* were detected between the genotypes suggesting potential disturbances in the regulation of apoptosis (Supplemental Fig. 4E). Furthermore, the basal epithelial cells and macrophages have been shown to act as phagocytes and clear away apoptotic cells in the HFs (Mesa *et al*. 2015), and induction of *Bnip3* has been associated to autophagy of the keratinocytes themselves (Moriyama et al. 2014), which may hamper detection of changes in apoptosis. Interestingly, it has also been shown that the TGFβ-induced regression phase reduces the SC pool (Mesa *et al*. 2015). TGFβ1 is also an important factor in the regulation of the HF bulge SC differentiation (Kopan and Weintraub 1993, Yamamoto *et al*. 2003) and has been shown to control mesenchymal SC differentiation to smooth muscle cells via regulation of the Jag1/Notch signaling in vascular development (Kurpinski *et al*. 2010). Furthermore, in line with our current findings, a recent study showed that endothelial cell specific *Hif-p4h-2* deletion in the lungs increases both TGFβ and NOTCH3 signaling due to HIF2α stabilization (Wang *et al*. 2016).

In conclusion, our results show that *Hif-p4h-2* expression in the mesenchymal *FoxD1*-lineage cells is fundamental for normal development of the truncal hair and HFs, and the lack of *Hif-p4h-2* in the *FoxD1*^*+*^ dermal fibroblasts and DP cells causes congenital alopecia. The most superficial parts of the skin have the most severe physiological hypoxia, as they are furthest away from the blood vessels that are located in the skin dermis and HIF has been reported to be endogenously stabilized in these parts of the skin (Cowburn *et al*. 2014, Evans *et al*. 2006, Rosenberger *et al*. 2007). As the HF morphogenesis proceeds, the dermal placode cells, that later develop into DP cells, travel closer to the hypodermal area, where there is a higher oxygen concentration and thus the endogenous HIF stabilization should normally be reduced or disappear in the DP cells. The highest oxygen concentration and hence lowest HIF amount in the DP cells should be when the HF is at its longest in the end of HF morphogenesis, before the beginning of the first HF cycle, or catagen. However, in our cKO mice HIF is constantly stabilized in the DP cells regardless of the oxygen concentration, which results in activation of the hypoxia response pathway and misregulation of Notch and TGFβ signaling (Supplemental Fig. 6). This leads to an imbalance in the reciprocal signaling between the DP cells and the bulge SCs causing reduced differentiation of the HF keratinocytes. Due to the lack of differentiation, HFs cannot maintain their proper structure, which results in HF cyst formation and poor hair production. In the poorly differentiated epidermal and HF cells keratin expression is significantly increased and the hair shaft production is disturbed resulting in alopecia in the mutant mice.

## MATERIALS AND METHODS

### Mice

In brief, a conditional *Hif-p4h-2* targeting construct resulting in the deletion of exon 3 encoding two catalytically critical residues, a histidine and an arginine required for binding of Fe^2+^ and 2-oxoglutarate (Epstein *et al*. 2001), respectively, was generated from a 7-kb genomic clone from Lambda FIX Library (Stratagene) (Supplemental Fig. S1SA). The targeting construct was electroporated into mouse embryonic stem (ES) cells and positively targeted ES cell clones were identified by Southern blotting (Supplemental Fig. S1B). Correctly targeted ES clones were used to generate chimeric mice via blastocyst injections in the Biocenter Oulu Transgenic Core Facility. The chimeras were crossed with C57/Bl6N mice and the offspring were genotyped by PCR (Supplemental Fig. S1C). To inactivate *Hif-p4h-2* in *FoxD1*-lineage cells, the *Hif-p4h-2*^*loxP/loxP*^ mice were bred with a *FoxD1-Cre* mouse line (Jackson B6; 129S4-Foxd1^tm1(GFP/cre)Amc^/J) (Humphreys *et al*. 2008) purchased from The Jackson Laboratory. Deletion of exon 3 and reduced amount of HIF-P4H-2 were confirmed by PCR and Western blotting (Supplemental Fig. S1D-E). Both female and male mice were used in the study since they expressed the same phenotype. The double *Cre*-reporter Rosa26^*mT/mG*^ mice (The Jackson Laboratory, 007676) (Muzumdar *et al*. 2007) were a kind gift from Prof. Seppo Vainio, University of Oulu. The Animal Experiment Board of Finland, following the regulations of the EU Directive 86/609/EEC, the European Convention ETS123, and the national legislation of Finland, approved the animal experiments in this study. Recommendations concerning laboratory animal experiments and handling given by the Federation of European Laboratory Animal Science Associations (FELASA), and the Finnish and EU legislations were followed.

### Tissue preparation and processing

Mouse skin biopsies were taken as indicated (Fig. 1A). Samples were fixed overnight in 10% phosphate buffered formalin and embedded in paraffin. 5-µm sections were cut with Thermo Scientific Microm HM355S Microtome. For cryosections, cranial skin biopsies were prepared from the cranial area (frontal to occipital area) at P21 and dorsal skin biopsies were prepared from P14, P21 and P27 mice. In brief, the tissue was coated with Tissue tek® O.C.T. embedding compound (Sakura, SA62550), immediately immersed in liquid nitrogen and placed in a Tissue tek® Cryomold (Sakura, SA62534). Cryoblocks were cut with Cryotome Leica CM3050S.

### Immunohistochemistry, immunofluorescence and morphometric analysis

Paraffin sections were stained with hematoxylin and eosin. For immunohistochemical and immunofluorescent stainings, the samples were pre-treated with citric acid before staining with antibodies shown in Supplemental Table 1. Apoptosis was analyzed using TUNEL assay by an *In Situ* Cell Death Detection kit (11684795910, Roche). As counterstain for IF and IHC stainings Hoechst or DAPI staining was used. For the analysis of *Rosa26*^*mT/mG*^ expression, *Rosa26*^*mTmG*^; *FoxD1*^*cre/+*^ dorsal skin cryosections were air-dried for 1 h, fixed with 10% phosphate buffered formalin for 10 min, washed with isopropanol and counterstained with Hoechst diluted in isopropanol, and mounted. Morphometric analyses were performed with Adobe Photoshop CS software from 4-18 sections/mouse and from 4-35 mice per genotype as indicated in the figure legends. Time points included in the analyses ranged from E18.5 to P29. In PCNA and TUNEL morphometric analyses the HS, keratin mass inside the cysts and sebaceous glands were not calculated to the HF area. The number of pSMAD2^+^ HF keratinocytes per total DAPI^+^ cell number was calculated. The visual field analyzed was a 10x magnification field.

### Western blotting

Protein lysates were prepared from snap-frozen dorsal skin with urea buffer (8 M Urea, 40 mM Tris-HCl, 2.5 mM EDTA, pH 8.0) containing phosphatase (PhosSTOP, Roche) and protease inhibitors (cOmplete Protease Inhibitor Cocktail Tablet EDTA-free, Roche). The lysates were analyzed by Western blotting with antibodies shown in Supplemental Table 1.

### Quantitative RT-PCR (qPCR)

Total RNA was isolated from dorsal skin using TriPure Isolation Reagent (11667157001, Roche) and treated with DNase I (#EN0521, Thermo Fisher Scientific). Reverse transcription of 1 μg RNA/20μl was performed with the iScript™ cDNA Synthesis Kit (1708890, Bio-Rad). qPCR was performed with iTaq™ Universal SYBR^®^ Green Supermix (1725120, Biorad). To minimize the variation of the housekeeping genes, we used the geometrical mean of β-actin and GAPDH for normalization of the data (Vandesompele *et al*. 2002). The data values are shown as relative levels normalized to the E18.5 control expression levels. qPCR primers are shown in Supplemental Table 2. The number of samples analyzed at different time points was: E18.5 control/cKO (n=6/6); P4 (n=5/5); P8 (n=4/4); P14 (n=6/6); P16 (n=5/5); P18 (n=5/4); P21 (n=7/5); and P24 (n=4/7).

### Electron microscopy

Transmission electron microscopy (TEM) and scanning electron microscopy (SEM) were performed in the Biocenter Oulu Electron Microscopy Core Facility. For TEM analysis P14 and P24 dorsal skin biopsies were processed and analyzed as described in Kutchuk *et al*. 2015.

For SEM analysis P24 dorsal skin biopsies were fixed in 2.5% glutaraldehyde in 0.1 M phosphate buffer, dehydrated in graded ethanol series and dried using critical point dryer (K850, Quorum Technologies). Dried samples were attached to aluminium specimen mount using double sided carbon tape and coated with 5 nm of platinum (Q150T ES, Quorum Technologies). Samples were examined in Σigma HD VP SEM (Carl Zeiss Microscopy).

### Statistical analyses

The statistical analyses were performed using Student’s *t* test or Welch’s *t* test. The data are shown as the means ± S.D. Values of p < 0.05 were considered statistically significant, * p < 0.05, ** p < 0.01, *** p < 0.001).

## ACKNOWLEDGEMENTS

We thank Prof. Seppo Vainio for the *Rosa26*^*mT/mG*^ mice, PhD Ari-Pekka Kvist for help in statistical analysis, Minna Siurua and Raija Salmu for expert technical assistance, Biocenter Oulu transgenic animal and electron microscopy core facilities supported by University of Oulu and Biocenter Finland, and the Laboratory Animal Center of University of Oulu. This study was supported by the Academy of Finland Center of Excellence 2012-2017 Grant 251314 (JM) and Academy Project Grant 296498 (JM), S. Jusélius Foundation (JM), Jane and Aatos Erkko Foundation (JM), FibroGen, Inc. (JM), The Finnish Cultural Foundation (A-HR) and the Instrumentarium Science Foundation (A-HR).

## AUTHORSHIP CONTRIBUTION

JM and JMM supervised and designed the study. JMM generated the *Hif-p4h-2* conditional mouse line. A-HR and MM carried out most of the experiments. A-HR wrote the manuscript with JMM and JM. AL and AR contributed to preliminary experiments. IM analyzed the TEM and SEM samples with A-HR. and JMM. RH contributed in designing the study.

## REFERENCES

Alam H, Sehgal L, Kundu ST, Dalal SN, Vaidya MM. 2011. Novel function of keratins 5 and 14 in proliferation and differentiation of stratified epithelial cells. Mol Biol Cell 22: 4068–4078.

Alibardi L, Bernd N. 2013. Immunolocalization of junctional proteins in human hairs indicates that the membrane complex stabilizes the inner root sheath while desmosomes contact the companion layer through specific keratins. Acta Histochem 115: 519–526.

Alonso L, Fuchs E. 2006. The hair cycle. J Cell Sci 119: 391–393.

Andersson ER, Sandberg R, Lendahl U. 2011. Notch signaling: simplicity in design, versatility in function. Development 138: 3593–3612.

Berra E, Benizri E, Ginouvès A, Volmat V, Roux D, Pouysségur J. 2003. HIF prolyl-hydroxylase 2 is the key oxygen sensor setting low steady-state levels of HIF-1α in normoxia. EMBO J 22: 4082–4090.

Borggrefe T, Lauth M, Zwijsen A, Huylebroeck D, Oswald F, Giaimo BD. 2016. The Notch intracellular domain integrates signals from Wnt, Hedgehog, TGFβ/BMP and hypoxia pathways. Biochim Biophys Acta 1863: 303–313.

Bose A, Teh MT, Mackenzie IC, Waseem A. 2013. Keratin k15 as a biomarker of epidermal stem cells. Int J Mol Sci 14: 19385–19398.

Botchkarev VA, Paus R. 2003. Molecular biology of hair morphogenesis: development and cycling. J Exp Zool B Mol Dev Evol 298: 164–180.

Brou C, Logeat F, Gupta N, Bessia C, LeBail O, Doedens JR, Cumano A, Roux P, Black RA, Israël A. 2000. A novel proteolytic cleavage involved in Notch signaling: the role of the disintegrin-metalloprotease TACE. Mol Cell 5: 207–216.

Bruick RK, McKnight SL. 2001. A conserved family of prolyl-4-hydroxylases that modify HIF. Science 294: 1337–1340.

Burton TR, Gibson SB. 2009. The role of Bcl-2 family member BNIP3 in cell death and disease: NIPping at the heels of cell death. Cell Death Differ 16: 515–523.

Cotsarelis G, Sun TT, Lavker RM. 1990. Label-retaining cells reside in the bulge area of pilosebaceous unit: implications for follicular stem cells, hair cycle, and skin carcinogenesis. Cell 61: 1329–1337.

Couly G, Le Douarin NM. 1988. The fate map of the cephalic neural primordium at the presomitic to the 3-somite stage in the avian embryo. Development 103 Suppl: 101–113.

Cowburn AS, Alexander LEC, Southwood M, Nizet V, Chilvers ER, Johnson RS. 2014. Epidermal deletion of HIF-2α stimulates wound closure. J Invest Dermatol 134: 801–808.

Driskell RR, Clavel C, Rendl M, Watt FM. 2011. Hair follicle dermal papilla cells at a glance. J Cell Sci 124: 1179–1182.

Dunwoodie SL. 2009. The role of hypoxia in development of the mammalian embryo. Dev Cell 17: 755–773.

Epstein AC, Gleadle JM, McNeill LA, Hewitson KS, O’Rourke J, Mole DR, Mukherji M, Metzen E, Wilson MI, Dhanda A et al. 2001. C. elegans EGL-9 and mammalian homologs define a family of dioxygenases that regulate HIF by prolyl hydroxylation. Cell 107:43–54.

Evans SM, Schrlau AE, Chalian AA, Zhang P, Koch CJ. 2006. Oxygen levels in normal and previously irradiated human skin as assessed by EF5 binding. J Invest Dermatol 126: 2596–2606.

Foitzik K, Paus R, Doetschman T, Dotto GP. 1999. The TGF-β2 isoform is both a required and sufficient inducer of murine hair follicle morphogenesis. Dev Biol 212: 278–289.

Fuchs E. 2007. Scratching the surface of skin development. Nature 445: 834–842.

Giaccia AJ, Simon MC, Johnson R. 2004. The biology of hypoxia: the role of oxygen sensing in development, normal function, and disease. Genes Dev 18: 2183–2194.

Gordan JD, Simon MC. 2007. Hypoxia-inducible factors: central regulators of the tumor phenotype. Curr Opin Genet Dev 17: 71–77.

Guo L, Yu QC, Fuchs E. 1993. Targeting expression of keratinocyte growth factor to keratinocytes elicits striking changes in epithelial differentiation in transgenic mice. EMBO J 12: 973–986.

Gustafsson MV, Zheng X, Pereira T, Gradin K, Jin S, Lundkvist J, Ruas JL, Poellinger L, Lendahl U, Bondesson M. 2005. Hypoxia requires notch signaling to maintain the undifferentiated cell state. Dev Cell 9: 617–628.

Higgins CA, Westgate GE, Jahoda CA. 2009. From telogen to exogen: mechanisms underlying formation and subsequent loss of the hair club fiber. J Invest Dermatol 129: 2100–2108.

Hsu YC, Li L, Fuchs E. 2014a. Emerging interactions between skin stem cells and their niches. Nat Med 20: 847–856.

Hsu YC, Li L, Fuchs E. 2014b. Transit-amplifying cells orchestrate stem cell activity and tissue regeneration. Cell 157: 935–949.

Hu YY, Fu LA, Li SZ, Chen Y, Li JC, Han J, Liang L, Li L, Ji CC, Zheng MH et al. 2014. Hif-1α and Hif-2α differentially regulate Notch signaling through competitive interaction with the intracellular domain of Notch receptors in glioma stem cells. Cancer Lett 349: 67–76.

Humphreys BD, Valerius MT, Kobayashi A, Mugford JW, Soeung S, Duffield JS, McMahon AP, Bonventre JV. 2008. Intrinsic epithelial cells repair the kidney after injury. Cell Stem Cell 2: 284–291.

Imamura Y, Tomita S, Imanishi M, Kihira Y, Ikeda Y, Ishizawa K, Tsuchiya K, Tamaki T. 2014. HIF-2α/ARNT complex regulates hair development via induction of p21(Waf1/Cip1) and p27(Kip1). FASEB J 28: 2517–2524.

Ivan M, Haberberger T, Gervasi DC, Michelson KS, Günzler V, Kondo K, Yang H, Sorokina I, Conaway RC, Conaway JW et al. 2002. Biochemical purification and pharmacological inhibition of a mammalian prolyl hydroxylase acting on hypoxia-inducible factor. Proc Natl Acad Sci USA 99: 13459–13464.

Ivan M, Kondo K, Yang H, Kim W, Valiando J, Ohh M, Salic A, Asara JM, Lane WS, Kaelin WG Jr. 2001. HIFα targeted for VHL-mediated destruction by proline hydroxylation: implications for O_2_sensing. Science 292: 464–468.

Jaakkola P, Mole DR, Tian YM, Wilson MI, Gielbert J, Gaskell SJ, von Kriegsheim A, Hebestreit HF, Mukherji M, Schofield CJ et al. 2001. Targeting of HIF-α to the von Hippel-Lindau ubiquitylation complex by O_2_-regulated prolyl hydroxylation. Science 292: 468–472.

Jamora C, Lee P, Kocieniewski P, Azhar M, Hosokawa R, Chai Y, Fuchs E. 2005. A signaling pathway involving TGF-β2 and snail in hair follicle morphogenesis. PLoS Biol 3: e11.

Joshi RS. 2011. The inner root sheath and the men associated with it eponymically. Int J Trichology 3: 57–62.

Kobayashi H, Liu Q, Binns TC, Urrutia AA, Davidoff O, Kapitsinou PP, Pfaff AS, Olauson H, Wernerson A, Fogo AB et al. 2016. Distinct subpopulations of FOXD1 stroma-derived cells regulate renal erythropoietin. J Clin Invest 126: 1926–1938.

Kopan R, Weintraub H. 1993. Mouse notch: expression in hair follicles correlates with cell fate determination. J Cell Biol 121: 631–641.

Kothari S, Cizeau J, McMillan-Ward E, Israels SJ, Bailes M, Ens K, Kirshenbaum LA, Gibson SB. 2003. BNIP3 plays a role in hypoxic cell death in human epithelial cells that is inhibited by growth factors EGF and IGF. Oncogene 22: 4734–4744.

Krause K, Foitzik K. 2006. Biology of the hair follicle: the basics. Semin Cutan Med Surg 25: 2–10.

Kurpinski K, Lam H, Chu J, Wang A, Kim A, Tsay E, Agrawal S, Schaffer DV, Li S. 2010. Transforming growth factor-β and notch signaling mediate stem cell differentiation into smooth muscle cells. Stem Cells 28: 734–742.

Kutchuk L, Laitala A, Soueid-Bomgarten S, Shentzer P, Rosendahl AH, Eilot S, Grossman M, Sagi I, Sormunen R, Myllyharju J et al. 2015. Muscle composition is regulated by a Lox-TGFβ feedback loop. Development 142: 983–993.

Magerl M, Tobin DJ, Müller-Röver S, Hagen E, Lindner G, McKay IA, Paus R. 2001. Patterns of proliferation and apoptosis during murine hair follicle morphogenesis. J Invest Dermatol 116: 947–955.

Mesa KR, Rompolas P, Zito G, Myung P, Sun TY, Brown S, Gonzalez DG, Blagoev KB, Haberman AM, Greco V. 2015. Niche-induced cell death and epithelial phagocytosis regulate hair follicle stem cell pool. Nature 522: 94–97.

Minamishima YA, Moslehi J, Bardeesy N, Cullen D, Bronson RT, Kaelin WG,Jr. 2008. Somatic inactivation of the PHD2 prolyl hydroxylase causes polycythemia and congestive heart failure. Blood 111: 3236–3244.

Moll R, Divo M, Langbein L. 2008. The human keratins: biology and pathology. Histochem Cell Biol 129: 705–33.

Moriyama M, Moriyama H, Uda J, Matsuyama A, Osawa M, Hayakawa T. 2014 BNIP3 plays crucial roles in the differentiation and maintenance of epidermal keratinocytes. J Invest Dermatol 134:1627–1635.

Muzumdar MD, Tasic B, Miyamichi K, Li L, Luo L. 2007. A global double-fluorescent Cre reporter mouse. Genesis 45: 593–605.

Myllyharju J. 2013. Prolyl 4-hydroxylases, master regulators of the hypoxia response. Acta Physiol (Oxf) 208: 148–165.

Nowell C, Radtke F. 2013. Cutaneous Notch signaling in health and disease. Cold Spring Harb Perspect Med 3: a017772.

Olivera-Martinez I, Thelu J, Dhouailly D. 2004. Molecular mechanisms controlling dorsal dermis generation from the somitic dermomyotome. Int J Dev Biol 48: 93–101.

Pan Y, Lin MH, Tian X, Cheng HT, Gridley T, Shen J, Kopan R. 2004. γ-secretase functions through Notch signaling to maintain skin appendages but is not required for their patterning or initial morphogenesis. Dev Cell 7: 731–743.

Paus R, Müller-Röver S, Van Der Veen C, Maurer M, Eichmüller S, Ling G, Hofmann U, Foitzik K, Mecklenburg L, Handjiski B. 1999. A comprehensive guide for the recognition and classification of distinct stages of hair follicle morphogenesis. J Invest Dermatol 113: 523–532.

Rathman-Josserand M, Genty G, Lecardonnel J, Chabane S, Cousson A, Francois Michelet J, Bernard BA. 2013. Human hair follicle stem/progenitor cells express hypoxia markers. J Invest Dermatol 133: 2094–2097.

Rezvani HR, Ali N, Nissen LJ, Harfouche G, de Verneuil H, Taieb A, Mazurier F. 2011. HIF-1α in epidermis: oxygen sensing, cutaneous angiogenesis, cancer, and non-cancer disorders. J Invest Dermatol 131: 1793–1805.

Rosenberger C, Solovan C, Rosenberger AD, Jinping L, Treudler R, Frei U, Eckardt KU, Brown LF. 2007. Upregulation of hypoxia-inducible factors in normal and psoriatic skin. J Invest Dermatol 127: 2445–2452.

Roth KJ, Copple BL. 2015. Role of hypoxia-inducible factors in the development of liver fibrosis. Cell Mol Gastroenterol Hepatol 1: 589–597.

Roth W, Kumar V, Beer HD, Richter M, Wohlenberg C, Reuter U, Thiering S, Staratschek-Jox A, Hofmann A, Kreusch F et al. 2012. Keratin 1 maintains skin integrity and participates in an inflammatory network in skin through interleukin-18. J Cell Sci 125: 5269–5279.

Rundle CW, Bergman D, Goldenberg A, Jacob SE. 2017. Contact dermatitis considerations in atopic dermatitis. Clin Dermatol 35: 367–374.

Schlake T. 2007. Determination of hair structure and shape. Semin Cell Dev Biol 18: 267–273.

Schofield CJ, Ratcliffe PJ. 2004. Oxygen sensing by HIF hydroxylases. Nat Rev Mol Cell Biol 5: 343–354.

Soma T, Tsuji Y, Hibino T. 2002. Involvement of transforming growth factor-β2 in catagen induction during the human hair cycle. J Invest Dermatol 118: 993–997.

Stenn KS, Paus R. 2001. Controls of hair follicle cycling. Physiol Rev 81: 449–494.

Takeda K, Ho VC, Takeda H, Duan LJ, Nagy A, Fong GH. 2006. Placental but not heart defects are associated with elevated hypoxia-inducible factor α levels in mice lacking prolyl hydroxylase domain protein 2. Mol Cell Biol 26: 8336–8346.

Tsuji, K., Kitamura, S. and Makino, H. (2014). Hypoxia-Inducible Factor 1α regulates Branching Morphogenesis during Kidney Development. Biochem. Biophys. Res. Commun. 447, 108–114.

Ueno M, Maeno T, Nomura M, Aoyagi-Ikeda K, Matsui H, Hara K, Tanaka T, Iso T, Suga T, Kurabayashi M. 2011. Hypoxia-inducible factor-1α mediates TGF-β-induced PAI-1 production in alveolar macrophages in pulmonary fibrosis. Am J Physiol Lung Cell Mol Physiol 300: L740–52.

Vandesompele J, De Preter K, Pattyn F, Poppe B, Van Roy N, De Paepe A, Speleman F. 2002. Accurate normalization of real-time quantitative RT-PCR data by geometric averaging of multiple internal control genes. Genome Biol 3: RESEARCH0034.

Vauclair S, Nicolas M, Barrandon Y, Radtke F. 2005. Notch1 is essential for postnatal hair follicle development and homeostasis. Dev Biol 284: 184–193.

Wang S, Zeng H, Xie XJ, Tao YK, He X, Roman RJ, Aschner JL, Chen JX. 2016. Loss of prolyl hydroxylase domain protein 2 in vascular endothelium increases pericyte coverage and promotes pulmonary arterial remodeling. Oncotarget 7: 58848–58861.

Watt FM, Estrach S, Ambler CA. 2008. Epidermal Notch signalling: differentiation, cancer and adhesion. Curr Opin Cell Biol 20: 171–179.

Weber S, Niessen MT, Prox J, Lullmann-Rauch R, Schmitz A, Schwanbeck R, Blobel CP, Jorissen E, de Strooper B, Niessen CM et al. 2011. The disintegrin/metalloproteinase Adam10 is essential for epidermal integrity and Notch-mediated signaling. Development 138: 495–505.

Yamamoto N, Tanigaki K, Han H, Hiai H, Honjo T. 2003. Notch/RBP-J signaling regulates epidermis/hair fate determination of hair follicular stem cells. Curr Biol 13: 333–338.

Zhao B, Guan H, Liu JQ, Zheng Z, Zhou Q, Zhang J, Su LL, Hu DH. 2017. Hypoxia drives the transition of human dermal fibroblasts to a myofibroblast-like phenotype via the TGF-β1/Smad3 pathway. Int J Mol Med 39: 153–159.

Zheng X, Narayanan S, Zheng X, Luecke-Johansson S, Gradin K, Catrina SB, Poellinger L, Pereira TS. 2017. A Notch-independent mechanism contributes to the induction of Hes1 gene expression in response to hypoxia in P19 cells. Exp Cell Res 358: 129–139.

