## Supplemental Figures for "Deletion of hypoxia-inducible factor prolyl 4-hydroxylase 2 in *FoxD1*-lineage mesenchymal cells leads to congenital truncal alopecia"

Rosendahl_Supplemental Fig. S1

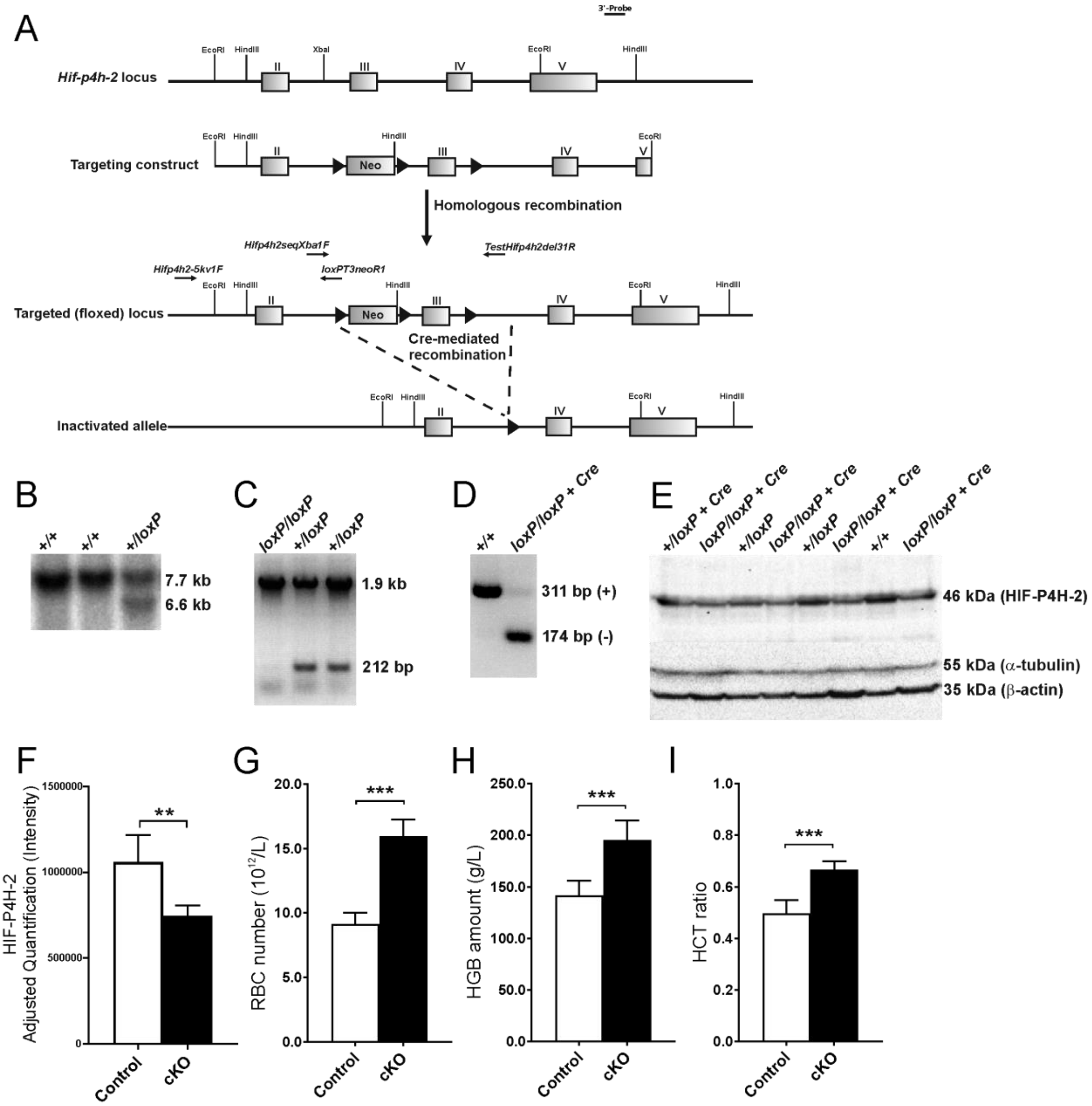

**Supplemental Figure S1.** Generation of *Hif-p4h-2*$^{loxP/loxP}$*;FoxD1*$^{Cre/+}$ (cKO) knock-out mice. (A) Schematic representation of the gene targeting strategy. The organization of the wild-type (wt) *Hif-p4h-2* gene, the targeting construct, and the structure of the locus following gene targeting are shown. Exons are depicted as gray boxes and numbered. *LoxP*-sequences are depicted as black arrowheads. The locations of PCR primers, Hind-III recognition sites and the 3'probe used for genotyping are shown in the picture. (B) Targeting was identified in ES cell lines by using the 3' probe and Hind-III digestion in Southern hybridization, where 7.7-kb wt (+) and 6.6-kb *Hif-p4h-2* conditional alleles (*loxP*) were identified, as expected. (C) Genotyping by PCR using the primer pairs Hifp4h2seqXba1F/TestHifp4h2del31R and Hifp4h2-5kv1F/loxPT3neoR1 from the F2 generation showed 212-bp and 1.9-kb bands for wt (+) and *Hif-p4h-2* conditional (*loxP/loxP*) alleles, respectively. (D) *FoxD1Cre*-mediated deletion of the *Hif-p4h-2* exon 3 was confirmed at the mRNA level from kidney by PCR primers mHifp4h2ex2F and mHifp4h2ex4R from exons 2 and 4, respectively, resulting in 311-bp wt (+) and 174-bp mutant (-) bands, as expected. (E) Western blotting and (F) quantification of HIF-P4H-2 from control and *Hif-p4h-2*$^{loxP/loxP}$*;FoxD1*$^{Cre/+}$ skin. (G-I) Blood analysis of the control and cKO mice showed increased levels of red blood cells (RBC), hemoglobin (HGB) and hematocrit (HCT).

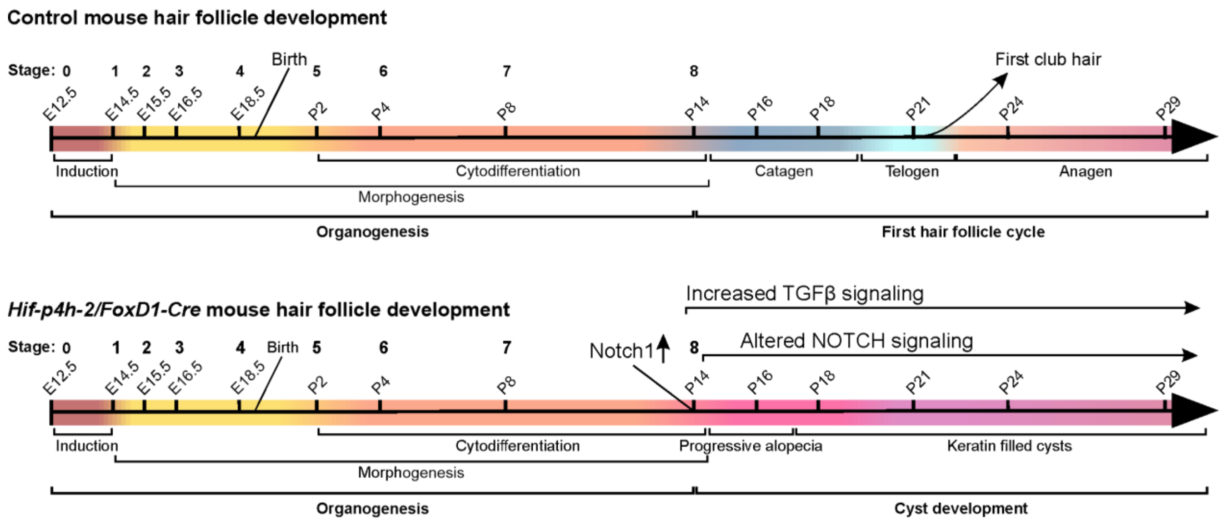


**Supplemental Figure S2.** Schematic represention of the timeline of HF development in control and cKO mice. A progressive alopecia starts at P15 in the cKO mice and evolves to keratin-filled epidermal cysts. TGFβ signaling is upregulated and Notch signaling is altered starting from P14 in the cKO mice relative to the control.

Rosendahl_Supplemental Fig. S3

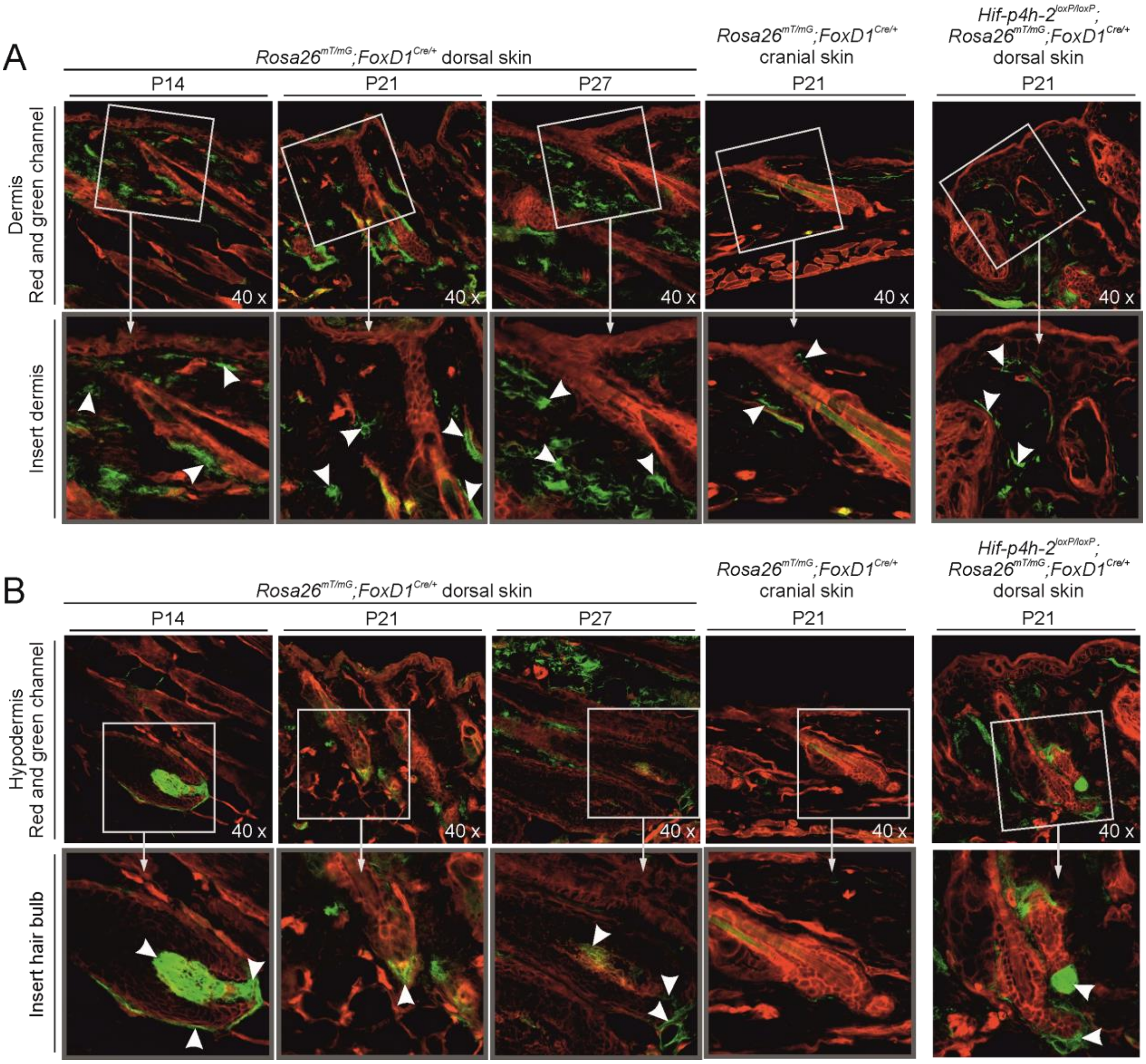


**Supplemental Figure S3.** Distribution of *FoxD1-Cre* expressing cells in mouse skin. (A) Dorsal dermis at P14 (morphogenesis, stage 8), P21 (telogen), and P27 (anagen) and cranial skin dermis at P21. (B) The dorsal hypodermis and the hair bulbs at P14, P21, and P27, and the hair bulb in cranial hypodermis at P21. Numerous green *FoxD1*$^+$ cells (indicated with arrowheads in the magnified inserts) are observed in the dorsal dermis, while they are rare in the cranial dermis.

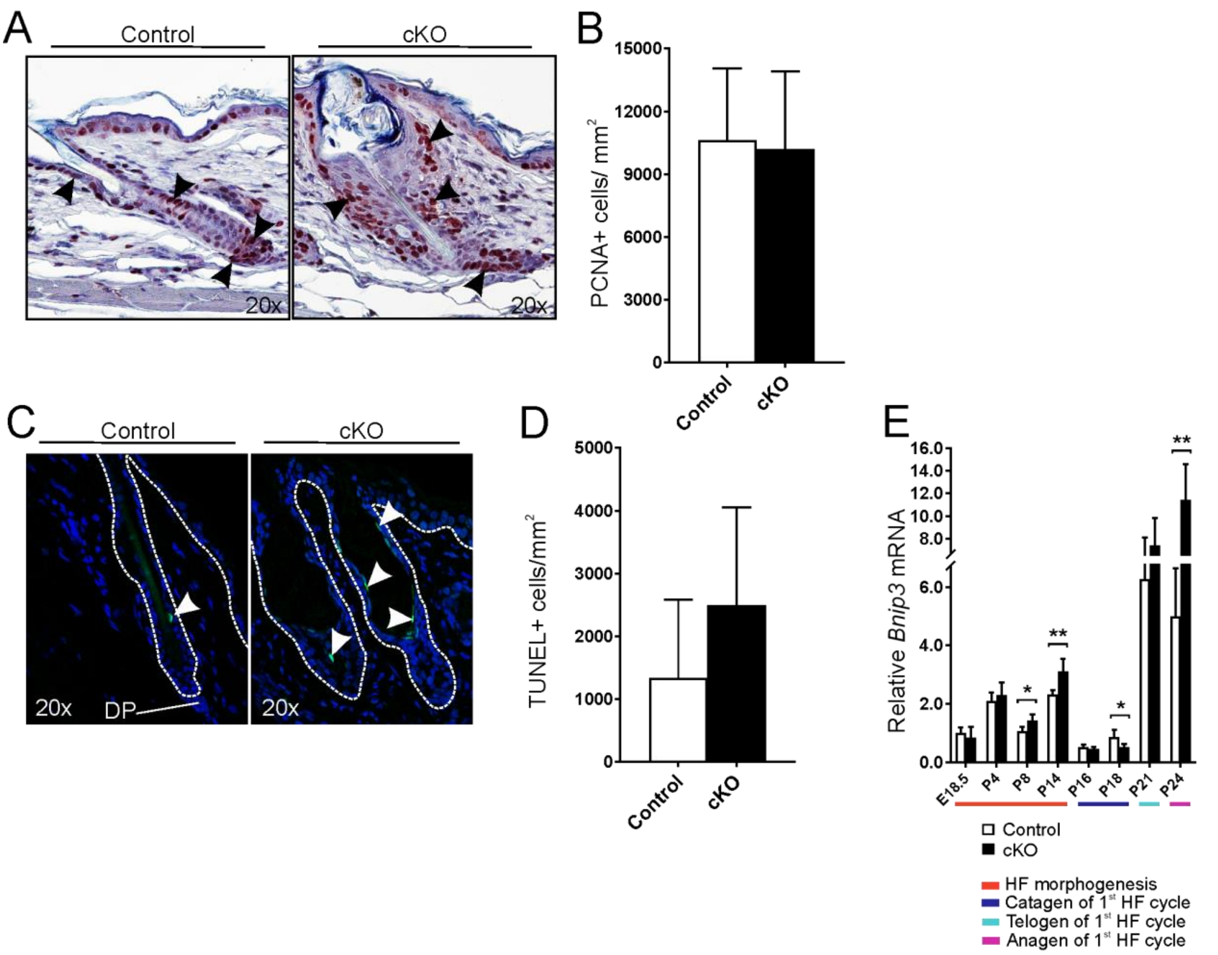


**Supplemental Figure S4.** Apoptosis and proliferation of cells is unaffected in the cKO mouse skin. (A) Analysis of proliferating cells (indicated by arrowheads) in the skin (P21) by immunohistochemical staining of PCNA. (B) Morphometric analysis of PCNA$^+$ cells in the skin (P21). Control (n = 5), cKO (n = 5). (C) Analysis of apoptotic cells (indicated by arrowheads) in the skin (P21) by TUNEL staining. (D) Morphometric analysis of apoptotic cells in the skin (P21). Control (n = 5), cKO (n = 5). (E) qPCR analysis of *Bnip3* mRNA expression at indicated time points. The colours beneath the bar charts indicate the HF cycle stages, n = 4-7 per genotype. Data are presented as mean ± S.D. * P<0.05; ** P<0.01.

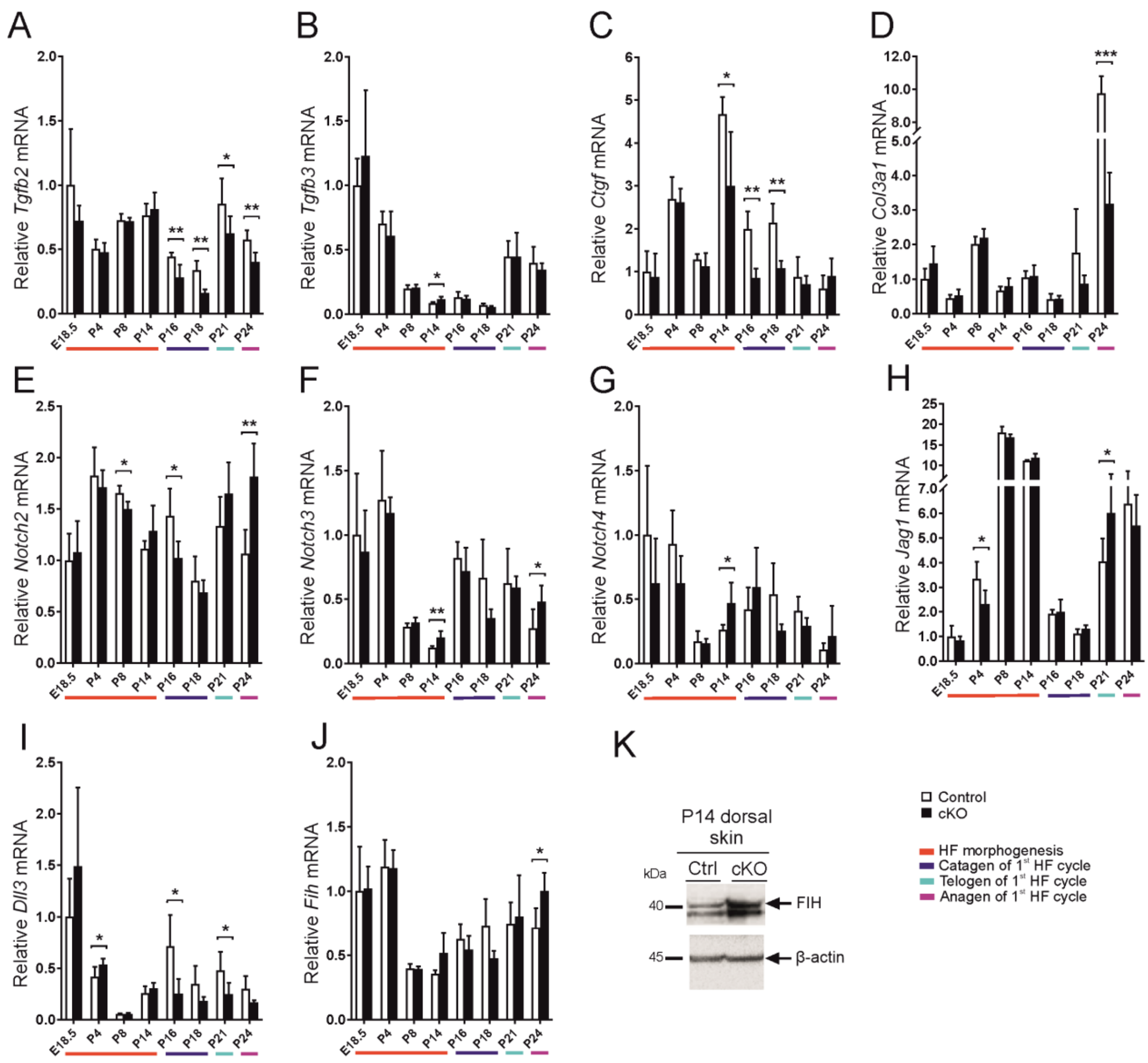


**Supplemental Figure S5.** Expression of TGFβ and Notch pathway related genes is altered in the cKO mice. qPCR analysis of the mRNA expression of the TGFβ isoforms *Tgfb2* (A) and *Tgfb3* (B); TGFβ target genes *Ctgf* (C) and *Col3a1* (D); NOTCH isoforms *Notch2* (E), *Notch3* (F) and *Notch4* (G); and NOTCH ligands *Jag1* (H) and *Dll3* (I); and *Fih* (J) at the indicated time points in the skin. The colours beneath the bar charts indicate the HF cycle stages, n = 4-7 per genotype. (K) Western blot analysis of FIH in P14 dorsal skin samples. Data are presented as mean ± S.D. * P<0.05; ** P<0.01; *** P<0.001.

Rosendahl_Supplemental Fig. S6

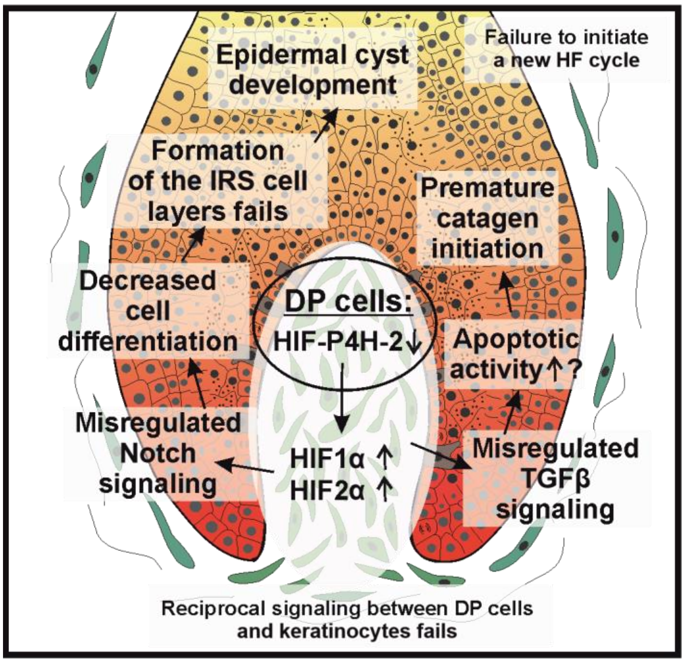


Cross section of hair follicle

**Supplemental Figure S6.** Schematic summary of the effects of HIF-P4H-2 depletion on HF development. As HIF-P4H-2 is lacking in the DP cells, HIF1 and 2 are stabilized and induce expression of HIF target genes. This also affects Notch and TGFβ signaling leading to disturbed homeostasis between the three signaling pathways. As a result, the receptor-ligand interactions between the DP cells and keratinocytes are altered, the cell differentiation fails, the formation of the supporting IRS layers fails and an epidermal cyst develops. Furthermore, apoptotic activity may be increased in the keratinocytes. Altogether, this leads to premature catagen initiation.
