## Supplemental Tables for "Deletion of hypoxia-inducible factor prolyl 4-hydroxylase 2 in *FoxD1*-lineage mesenchymal cells leads to congenital truncal alopecia"

**Table S1.** Antibodies and other reagents used for immunostaining or Western blot analyses.

| Name | Code | Company | Dilution |
|---|---|---|---|
| Adam10 | ab1997 | Abcam | WB 1:1000 |
| Activated Notch1 | ab8925 | Abcam | WB 1:500 |
| Cy3 | 715-165-150<br>711-165-152<br>712-165-150 | Jackson Immunoresearch | IF 1:300 |
| Keratin 1 | ab24643 | Abcam | IF 1:100 |
| Keratin 15 | ab2414 | Abcam | IF 1:100 |
| DAPI | D9542 | Sigma | IF 1:500 |
| FIH-1/HIF-1AN | NB100-428SS | Novus Biologicals | WB 1:500 |
| HDAC1 | #5356 | Cell Signaling Technology | WB 1:1000 |
| Hes1 | ab71559 | Abcam | WB 1:1000 |
| HIF1 alpha | NB100-479 | Novus Biologicals | WB 1:500 |
| HIF2 alpha | GTX30114 | GeneTex | WB 1:500 |
| Hoechst | 33258 | Sigma Aldrich | IF 1:10000 |
| Keratin 5 | PRB-160P | Biolegend | IF 1:100 |
| Loricrin | PRB-145P | Biolegend | IF 1:200 |
| Mac-3 | 550292 | BD Pharmingen™ | IHC 1:100 |
| Notch1 | ab27526 | Abcam | WB 1:500 |
| PCNA | sc-56 | Santa Cruz Biotechnology | IHC 1:100 |
| Phospho-Smad2 (Ser465/467) | #3101 | Cell Signaling Technology | IF 1:100 |
| α-Tubulin | B-6199 | Sigma Aldrich | WB 1:20000 |

**Table S2.** PCR primers

| Name | Gene | Oligo direction | Oligo sequence |
|---|---|---|---|
| Adam8 | ADAM metallopeptidase domain 8 | Forward | CCCAAGACCCATAGTGAAACCAA |
| | | Reverse | CTTTGGGGCATAAACAGGAACTG |
| Adam10 | ADAM metallopeptidase domain 10 | Forward | AACATCAGCTTCATGGTGAAACG |
| | | Reverse | CCAGGAACTTCTCCACACCAATA |
| Adam17 | ADAM metallopeptidase domain 17 | Forward | GGTTCTAGCCCACATAGGAGATG |
| | | Reverse | CCTCCAAAGTGGCTCTACGTTAT |
| β-Actin | β-Actin | Forward | AGAGGGAAATCGTGCGTGAC |
| | | Reverse | CAATAGTGATGACCTGGCCGT |
| Bnip3 | BCL2/adenovirus E1B interacting protein 3 | Forward | GCTCCAAGAGTTCTCACTGTGAC |
| | | Reverse | GTTTTTCTCGCCAAAGCTGTGGC |
| Col3a1 | Collagen type III alpha 1 chain | Forward | CTGTAACATGGAAACTGGGGAAA |
| | | Reverse | CCATAGCTGAACTGAAAACCACC |
| Cre | Cre recombinase (used for genotyping) | Forward | GCACGTTCACCGCATCAAC |
| | | Reverse | CGATGCAACGAGTGATGAGGTTC |
| Ctgf | Connective tissue growth factor | Forward | GGGCCTCTTCTGCGATTTC |
| | | Reverse | ATCCAGGCAAGTGCATTGGTA |
| Dll1 | Delta like canonical Notch ligand 1 | Forward | CAGGACCTTCTTTCGCGTATG |
| | | Reverse | AAGGGGAATCGGATGGGGTT |
| Dll3 | Delta like canonical Notch ligand 3 | Forward | GTCATACCAGCCCCTTCCATTTA |
| | | Reverse | GCGATGATAGAGAAGGGACAAGA |
| Dll4 | Delta like canonical Notch ligand 4 | Forward | TCCCAGGGACTCTATGTACCAAT |
| | | Reverse | CTGAGTAGGCTCCTGCCTTATAC |
| Eln | Elastin | Forward | TGTCCCACTGGGTTATCCCAT |
| | | Reverse | CAGCTACTCCATAGGGCAATTTC |
| Eno1 | Enolase 1 | Forward, reverse | QuantiTect Mm_Eno1_1_SG (QT00260442) |
| Fgf7 (Kgf) | Keratinocyte growth factor | Forward | AGCTGTTCCAAACAGAACAAAAGT |
| | | Reverse | ACCAATAACACGATTCCTCCTTCA |
| Fih | Factor inhibiting Hif | Forward | ATGAGTCCCAGCTACGAAGTTAC |
| | | Reverse | CAGTGCAGGATACACAAGGTTTG |
| Flg | Filaggrin | Forward | AGATGTCCGCTCTCCTGGAA |
| | | Reverse | TGGATTCTTCAAGACTGCCTGTA |
| Fn1 | Fibronectin | Forward, reverse | QuantiTect Mm_Fn1_1_SG (QT00135758) |
| GAPDH | Glyceraldehyde-3-phosphate dehydrogenase | Forward | TGTGTCCGTCGTGGATCTGA |
| | | Reverse | TTGCTGTTGAAGTCGCAGGAG |
| Hes1 | Hes family bHLH transcription factor 1 | Forward | CGGCATTCCAAGCTAGAGAAGG |
| | | Reverse | GGTAGGTCATGGCGTTGATCTG |
| Hes5 | Hes family bHLH transcription factor 5 | Forward | CATCAACAGCAGCATAGAGCAG |
| | | Reverse | GCGAAGGCTTTGCTGTGTTTCA |
| Hey1 | Hairy/enhancer-of-split related with YRPW motif 1 | Forward | TGAGCTGAGAAGGCTGGTAC |
| | | Reverse | ACCCCAAACTCCGATAGTCC |

**Table S2.** PCR primers continued

| Name | Gene | Oligo direction | Oligo sequence |
|---|---|---|---|
| Hey2 | Hairy/enhancer-of-split related with YRPW motif 2 | Forward | GACTTCATGAGCATTGGATTCCG |
| | | Reverse | CAGGTGCTGAGATGAGAGACAAG |
| HeyL | Hairy/enhancer-of-split related with YRPW motif-like | Forward | TGGGTCAAGAGAACGATCTTAGC |
| | | Reverse | CTATGATCCCTCTGCGCTTCTTC |
| Hif-p4h-1 | HIF prolyl 4-hydroxylase 1 | Forward | AGAACTGGGATGTTAAGGTGCAT |
| | | Reverse | GAAAATGAGCAACCGGTCAAAGAG |
| Hif-p4h-2 | HIF prolyl 4-hydroxylase 2 | Forward | GAGATGGAAGATGCGTGACA |
| | | Reverse | TTGCCTTCTGGAAAAATTCG |
| Hifp4h2-5KV1F | HIF prolyl 4-hydroxylase 2 (used for genotyping) | Forward | ACCAACCATTCCTAGAGAGTCATC |
| Hifp4h2seqXba1F | HIF prolyl 4-hydroxylase 2 (used for genotyping) | Forward | GCTCAGGAGCAGCAGTGTG |
| Hif-p4h-3 | HIF prolyl 4-hydroxylase 3 | Forward | CTTATTCAGGTAGTAGATACAGGTGATACA |
| | | Reverse | GCTGGGCAAATACTATGTCAAG |
| Hk2 | Hexokinase 2 | Forward, reverse | QuantiTect Mm_Hk2_1_SG (QT00155582) |
| Jag1 | Jagged 1 | Forward | CCTCGGGTCAGTTTGAGCTG |
| | | Reverse | CCTTGAGGCACACTTTGAAGTA |
| Jag2 | Jagged 2 | Forward | TGGCTGTCACCGAGGTCAA |
| | | Reverse | ACGTTCTTTCCTGCGCTTTC |
| Krt1 | Cytokeratin 1 | Forward | AGCTGAATCGAATGATCCAGAGA |
| | | Reverse | TGCTGTATTTGGGAGATCTGCTT |
| Krt5 | Cytokeratin 5 | Forward | GGTGGAGGACTACAAGAACAAGT |
| | | Reverse | GCATCCACATCCTTCTTCAACAT |
| Krt10 | Cytokeratin 10 | Forward | AGGACCAAGATACTAACAAAACCAA |
| | | Reverse | AGTGGCCCGTATGAAGAGACT |
| Krt14 | Cytokeratin 14 | Forward | CAAAGACTACAGCCCCTACTTCA |
| | | Reverse | CTCAAACTTGGTCCGGAAGTCAT |
| Krt15 | Cytokeratin 15 | Forward | GAGGTGAAGATCCGAGATTGGTA |
| | | Reverse | CCAGAATTTTGTCCCGGATCTCT |
| Ldha | Lactate dehydrogenase A | Forward, reverse | QuantiTect Mm_Ldha_1_SG (QT01045492) |
| Lor | Loricrin | Forward | CATGAATTTGCCTGAGGTTTCCA |
| | | Reverse | GGGAGGTAGTCATTCAGAAACCA |
| loxPT3neoR1 | Genotyping primer | Reverse | GCTATACGAAGTTATTAGGTCC |
| mHifp4h2ex2F | Genotyping primer | Forward | GCCATGGTTGCTTGTTACCC |
| mHifp4h2ex4R | Genotyping primer | Reverse | ACTTTAGCTCTCGCTCGCTC |
| Ngf | Nerve growth factor | Forward | GGGAGCGCATCGAGTTTTG |
| | | Reverse | CCAGTATAGAAAGCTGCGTCCTT |
| Notch1 | Notch 1 | Forward | GATGGCCTCAATGGGTACAAG |
| | | Reverse | TCGTTGTTGTTGATGTCACAGT |
| Notch2 | Notch 2 | Forward | GAGAAAAACCGCTGTCAGAATGG |
| | | Reverse | GGTGGAGTATTGGCAGTCCTC |

**Table S2.** PCR primers continued

| Name | Gene | Oligo direction | Oligo sequence |
|---|---|---|---|
| Notch3 | Notch 3 | Forward | AGATGCACTGGGAATGAAGAACA |
| | | Reverse | GCTCCTCTACCTTCAGTCTCTTG |
| Notch4 | Notch 4 | Forward | GAACGCGACATCAACGAGTG |
| | | Reverse | GGAACCCAAGGTGTTATGGCA |
| Pdk1 | Puryvate dehydrogenase kinase 1 | Forward, reverse | QuantiTect Mm_Pdk1_1_SG (QT00116396) |
| Postn | Periostin | Forward, reverse | QuantiTect Mm_Postn_1_SG (QT00150759) |
| Serpine1 (Pai-1) | Plasminogen activator inhibitor-1 | Forward, reverse | QuantiTect Mm_Serpine1_1_SG (QT00154756) |
| Slc2a1 | Glucose transporter 1 | Forward, reverse | QuantiTect Mm_Slc2a1_1_SG (QT01044953) |
| TestHifp4h2del31R | HIF prolyl 4-hydroxylase 2 (used for genotyping) | Reverse | CTGGCCCTTGTGTTTCCGAC |
| Tgfb1 | Transforming growth factor β1 | Forward | GAGCCCGAAGCGGACTACTA |
| | | Reverse | TGGTTTTCTCATAGATGGCGTTG |
| Tgfb2 | Transforming growth factor β2 | Forward | AGTTTACACTGCCCCTGCTG |
| | | Reverse | AGAGGTGCCATCAATACCTGC |
| Tgfb3 | Transforming growth factor β3 | Forward | CACCACAACCCACACCTGAT |
| | | Reverse | CAGGTTGCGGAAGCAGTAAT |
| Tnfa | Tumor necrosis factor α | Forward | GGTGCCTATGTCTCAGCCTCTT |
| | | Reverse | GCCATAGAACTGATGAGAGGGAG |
| Vegfa | Vascular endothelial growth factor A | Forward | CCACGTCAGAGAGCAACATCA |
| | | Reverse | TCATTCTCTCTATGTGCTGGCTTT |
